# A single cell atlas of human cornea that defines its development, limbal stem and progenitor cells and the interactions with the limbal niche

**DOI:** 10.1101/2020.07.09.195438

**Authors:** Joseph Collin, Rachel Queen, Darin Zerti, Sanja Bojic, Nicky Moyse, Marina Moya Molina, Chunbo Yang, Gary Reynolds, Rafiqul Hussain, Jonathan M Coxhead, Steven Lisgo, Deborah Henderson, Agatha Joseph, Paul Rooney, Saurabh Ghosh, Che Connon, Muzlifah Haniffa, Francisco Figueiredo, Lyle Armstrong, Majlinda Lako

## Abstract

To study the development and composition of human ocular surface, we performed single cell (sc) RNA-Seq at key embryonic, fetal and adult stages and generated the first atlas of the corneal cell types from development to adulthood. Our data indicate that during development, the conjunctival epithelium is the first to be specified from the ocular surface epithelium, followed by the corneal epithelium and the establishment of proliferative epithelial progenitors, which predate the formation of limbal niche by a few weeks. Bioinformatic comparison of adult cell clusters identified GPHA2, a novel cell-surface marker for quiescent limbal stem cells (qLSCs), whose function is to maintain qLSCs self-renewal. Combining scRNA- and ATAC-Seq analysis, we identified multiple upstream regulators for qLSCs and transit amplifying (TA) cells and demonstrated a close interaction between the immune cells and epithelial stem and progenitor cells in the cornea. RNA-Seq analysis indicated loss of qLSCs and acquisition of proliferative limbal basal epithelial progenitor markers during *ex vivo* limbal epithelial cell expansion, independently of the culture method used. Extending the single cell analyses to keratoconus, we were able to reveal activation of collagenase in the corneal stroma and a reduced pool of TA cells in the limbal epithelium as two key changes underlying the disease phenotype. Our scRNA- and ATAC-Seq data of developing and adult cornea in steady state and disease conditions provide a unique resource for defining pathways/genes that can lead to improvement in *ex vivo* expansion and differentiation methods for cell based replacement therapies and better understanding and treatment of ocular surface disorders.

**Key findings:**

- scRNA-Seq of adult human cornea and conjunctiva reveals the signature of various ocular surface cell populations
- scRNA-Seq of human developing cornea identifies stage-specific definitions of corneal epithelial, stromal and endothelial layers
- scRNA-Seq analysis results in identification of novel markers for qLSCs and TA cells
- Combined scRNA- and ATAC-Seq analysis reveals key transcriptional networks in qLSCs and TA cells and close interactions with immune cells
- Expansion of limbal epithelium results in downregulation of qLSCs and acquisition of proliferative limbal epithelial progenitor markers
- scRNA-Seq of keratoconus corneas reveals activation of collagenase in the corneal stroma and a reduced pool of TA cells in the limbal epithelium

**Graphical abstract:** Schematic presentation of main techniques and findings presented in this manuscript.

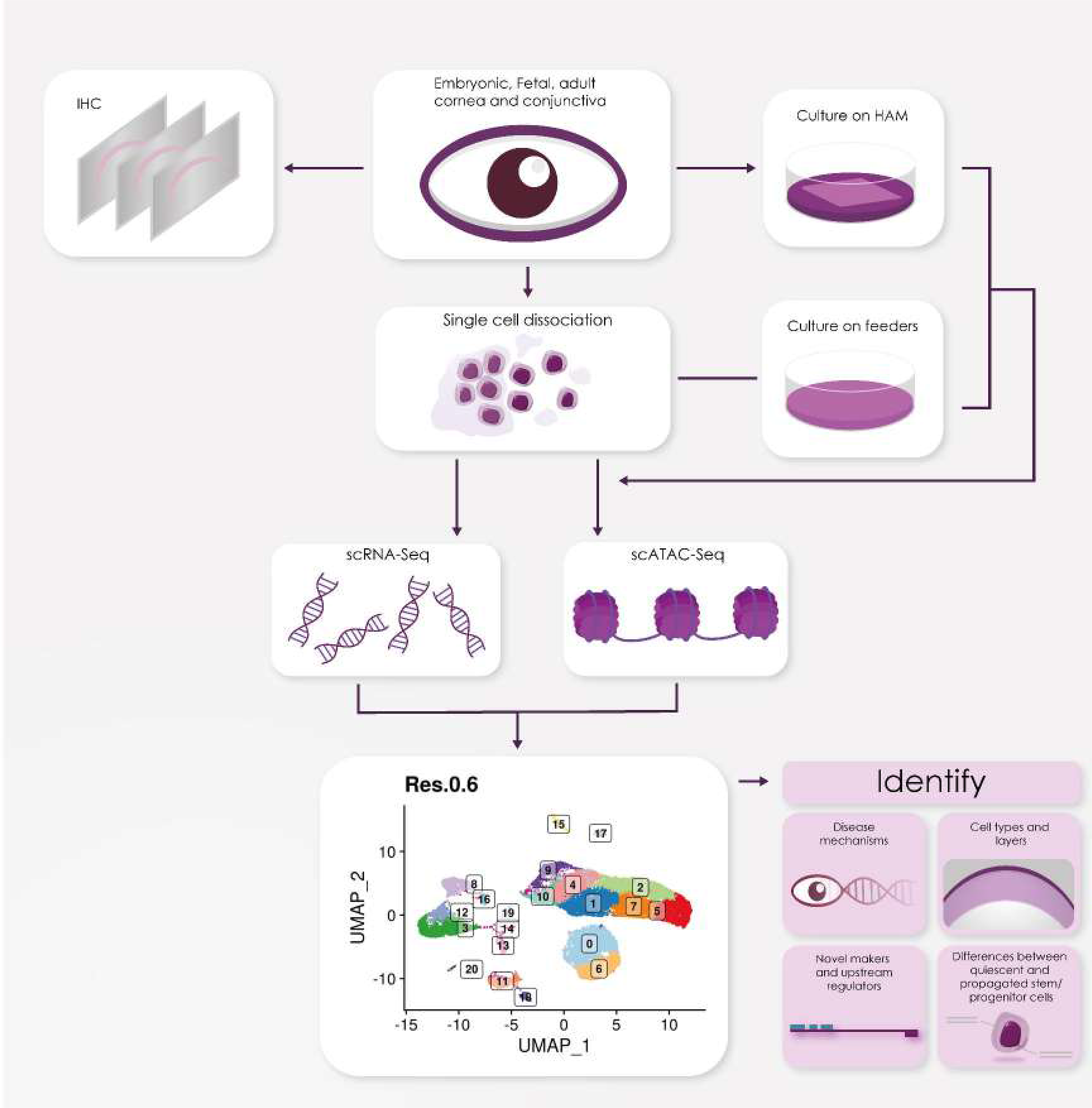

## Introduction

Cornea is the transparent front part of the eye, which together with the lens focuses the light onto retina for visual processing (Osei-Bempong et al., 2013). Corneal blindness is the 2^nd^ main cause of blindness worldwide accounting for 23 million people adding a huge burden to health care resources (Oliva et al., 2012; Pascolini and Mariotti, 2012; Whitcher et al., 2001). Often the only treatment is surgical transplantation of donor cornea, a therapeutic option that has been in practice for more than a century. In Europe, over 40,000 blind people are waiting for corneal transplant every year (Brown, 2010). This worldwide shortage of corneas results in about 10 million untreated patients globally and 1.5 million new cases of blindness annually (Whitcher et al., 2001).

Despite efforts to develop corneal substitutes, surgery with allogeneic donor tissue from cadavers has remained the gold standard for more than a century (Osei-Bempong et al., 2013). Transplantation of limbal stem cells (LSCs) and closely related epithelial cells have been used for corneal epithelial therapy (Kolli et al., 2009, 2014). However, these clinical techniques only address epithelial regeneration, and not restoration of a dysfunctional corneal stroma or endothelium. Clearly, there is an unmet need for the design of new smart biomaterials and stem cell therapies to create a whole cornea that is indistinguishable from the original native tissue and fulfill the natural functions of a transparent cornea.

The cornea is comprised of five layers: the outermost epithelium, Bowman’s layer, the stroma, the Descemet’s membrane and the endothelium (Sridhar, 2018). The stratified epithelium covers the outermost surface of the cornea and is divided from stroma by Bowman’s layer, a smooth, acellular layer made up of collagen fibrils and proteoglycans, which helps the cornea maintain its shape. There is high corneal epithelial cell turnover due to blinking as well as physical and chemical environmental insults. The renewal of corneal epithelium is sustained by the limbal stem cells (LSCs), which are located at the Palisades of Vogt at the limbal region that marks the transition zone between clear cornea and conjunctiva (Cotsarelis et al., 1989).

The corneal stroma occupies 90% of the corneal thickness (Hertsenberg and Funderburgh, 2015). The stroma is composed of water, proteoglycans and collagen fibrils, arranged in lamellae to reduce light scattering and enable corneal transparency. The stroma is populated by scarcely distributed keratocytes, which secrete the collagens and proteoglycans, in addition to a small population of corneal stromal stem cells (CSSCs), which are localized in the anterior peripheral (limbal) stroma near to LSCs. CSSCs display properties of mesenchymal stem cells, including clonal growth, multipotent differentiation, and expression of stem cell-specific markers and are characterized by the ability to divide extensively *in vitro* and to generate adult keratocytes (Pinnamaneni and Funderburgh, 2012). Genesis of corneal endothelium begins when periocular neural crest cells migrate between the presumptive corneal epithelium and lens vesicle and undergo a mesenchymal-to-endothelial transition to form a monolayer that occupies the posterior surface of the cornea (Lwigale, 2015). A major function of corneal endothelium is to maintain corneal transparency by regulating corneal hydration. The corneal endothelium comprises a single layer of closely interdigitating hexagonal cells, which secrete the Descemet membrane, a cell-free matrix that mostly consists of collagens. Unlike the corneal epithelial cells, endothelial cells in humans are not endogenously renewed or replaced during a lifetime and their cell density declines at an average of approximately 0.6% per year in normal corneas throughout human life.

By understanding the physical and cellular structure of cornea novel materials and new therapies for a number of ophthalmic indications can be designed. We performed single cell (sc) RNA-Seq of human cornea and surrounding conjunctiva during human development and in adulthood, in both steady state and disease conditions. Our data provide the first single cell atlas of developing and adult human cornea and surrounding conjunctiva that defines their development, limbal stem and progenitor cells and the interactions with the niche.

## Results

### scRNA-Seq of adult human cornea and surrounding conjunctiva reveals the presence of stem, progenitor and differentiated cells in the epithelial, stromal and endothelial layers

Human adult cornea and the surrounding conjunctiva were excised from four deceased donor eyes (51, 75, 83 and 86 years old) and dissociated to single cells. Approximately 10,000 cells were captured from each sample using the 10 X Chromium Single Cell 3’ Library & Gel Bead Kit Genomics (version 3). 21,343 cells were obtained from the four adult corneas after data integration and doublet cell exclusion. These were embedded using Uniform Manifold Approximation and Projection (UMAP) and clustered using Seurat graph based-clustering, which revealed the presence of 21 cell clusters (**Figure 1A**). Cluster identification was performed on the basis of marker genes (**Figure S1**, **Table S1**), bioinformatic data mining and immunohistochemical (IHC) analysis.

**Figure 1.**
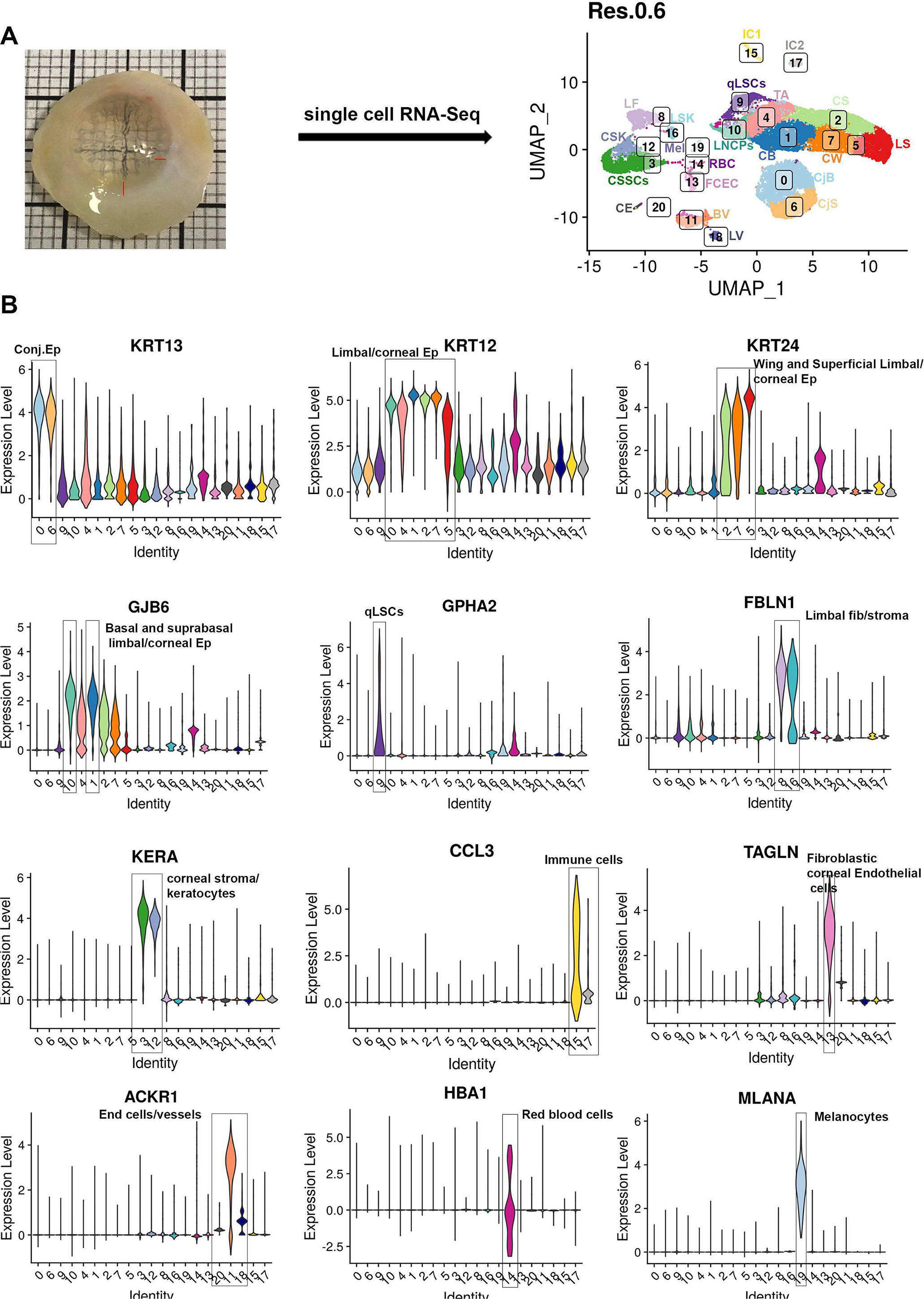
scRNA-Seq of adult human cornea and conjuctiva. (see also Figures S1-S14 and Table S1). **A**) UMAP of adult human cornea and conjunctiva showing the presence of stem, progenitor and differentiated cells in the epithelial, stromal and endothelial layers; **B**) Violin plots showing the presence of key markers for stem, progenitor and differentiated cells in the epithelial, stromal and endothelial compartments. *Abbreviations for panel 1A:* BV – blood vessels CB – corneal epithelial basal cells CS – corneal epithelial superficial cells CW – corneal wing cells CjB – conjunctival epithelial basal cells CjS – conjunctival epithelial superficial cells CE- corneal endothelium CSK – corneal stroma keratocytes CSSCs – corneal stromal stem cells FCEC – fibroblastic corneal endothelial cells IC1 – immune cells 1 IC2 – immune cells 2 LF – limbal fibroblasts LSK – limbal stroma keratocytes LNCPs – limbal neural crest progenitors qLSCs – quiescent limbal stem cells LS – limbal epithelial superficial cells LV – lymphatic vessels Mel – melanocytes RBC – red blood cells TA – transit amplifying cells *Abbreviations for panel 1B:* Conj- conjunctival Ep- epithelium Fib- fibroblasts End- endothelial

#### Identification of epithelial cell populations

Clusters 0 and 6 show high expression of Keratin 13 (*KRT13*) and 19 (*KRT19*) (Merjava et al., 2011; Ramirez-Miranda et al., 2011) as well as *S100A8* and *S100A9* (Li et al., 2011), which point to conjunctival cell fates (**Figure 1B** and **Figure S2A**). Differential gene expression analysis showed *KRT6A, KRT14* and *KRT15* to be more highly expressed in the cell cluster 0 (**Figure S2B**), suggestive of a basal conjunctival epithelium, which was confirmed by IHC (**Figure S2C**). Cluster 6 displayed higher expression of Mucin 4 (*MUC4*) and 1 (*MUC1*) as well as *KRT4*, indicative of a superficial conjunctival epithelium, which was also corroborated by IHC (**Figure S2C**).

Cell clusters 1, 2, 4, 5, 7 and 10 displayed high expression of keratin 12 (*KRT12*), whose expression is associated with corneal and limbal epithelium (Tanifuji-Terai et al., 2006) (**Figure 1B**). Of these, clusters 2, 5 and 7 displayed high expression of *KRT24*, whose expression is found in the suprabasal (also known as wing cells) and the superficial layers of corneal epithelium. Amphiregulin (*AREG*) was the most highly expressed gene in cluster 2 (**Figure S1, S3A**). Its expression was confined to the most superficial part of corneal epithelium, (**Figure S3B**), leading to definition of this cluster as corneal superficial epithelial cells. Differential gene expression analysis indicated *LYPD2* to be more highly expressed in cluster 5 (**Figure S3A**); hence, IHC with antibody raised against LYPD2 was carried out, showing a limbal superficial epithelial phenotype (**Figure S3C**). High expression of *KRT3* and *KRT12* with *KRT24* in cell cluster 7 indicates a central cornea wing or superficial epithelial cell fate. IHC analysis revealed KRT3 immunostaining in the corneal wing cells and its absence from the very flat squamous cells at the corneal superficial epithelium (**Figure S3D**); hence, this cluster was defined as corneal wing cells.

Cell clusters 1 shows high expression of gap junction protein beta 6 (*GJB6*), which is associated with a basal corneal epithelial phenotype (Shurman et al., 2005). *HES1* and *HES5*, whose expression is confined to the basal and immediate suprabasal corneal epithelium (Djalilian et al., 2008), were also highly expressed in cluster 1 (**Table S1**). IHC analysis revealed GJB6-immunostained cells to be located predominantly in the basal corneal epithelium (**Figure S4**), but not in the corneal superficial layers or the basal limbal epithelium, confirming the identity of cluster 1 as corneal basal epithelial cells.

Cluster 4, 9 and 10 were transcriptionally similar to each other (**Figure S1**), we thus performed a differential gene expression and violin plot analyses, which showed high expression of *KRT14* and *TP63* in all three clusters; however *KRT15*, a LSCs and progenitor marker (Chen et al., 2010; Yoshida et al., 2006) was highly expressed in clusters 9 and 4 but not in cluster 10 (**Figure S5A-D**). Through the differential gene expression analysis (**Figure S5A**), we were able to identify a unique marker for cluster 10, *CPVL*, which was not expressed in the other epithelial clusters (**Figure S6A**). CPVL immunopositive cells were found in the limbal stroma next to the limbal basal epithelial cells, with a small minority coexpressing TP63 (**Figure S6B-D**). This cluster also showed high expression of *PAX6* (**Table S1**), a transcription factor expressed in the limbal niche cells in the stroma (Chen et al., 2019), whose function is to maintain the phenotype of neural crest progenitors. In view of this, we speculated that cluster 10 may represent the limbal neural crest derived progenitors (LNCPs). IHC analysis revealed a large overlap in expression between the neural crest marker, MITF and the cluster 10 specific marker, CPVL (**Figure S6E, E’**), confirming the LNCPs nature of this cell population.

Cluster 4 showed high expression of LSCs and limbal progenitors (also known as transit amplifying (TA) cells)) markers including *KRT15, KRT14, CXCL14* (Ojeda et al., 2013) and *CDH13* (Mikhailova et al., 2015) (**Table S1**); however this cluster also expressed the tight junction transmembrane Claudin 1 (*CLDN1*) and 4 (*CLDN4*), which are present throughout the cell layers of corneal and conjunctival epithelium (Yoshida et al., 2009), suggesting that the cells in this cluster are probably distinct from the LSCs. A dual plot gene expression heatmap showed the highest co-expression of *KRT15* and *CLDN4* in cell cluster 4 and to a lesser extent in cluster 0 (**Figure S7A**). IHC with both antibodies showed the highest coexpression of these two markers in cells that are moving away from the basal layers of the limbal crypt, suggesting that these cells represent the TA cells, which migrate from the limbus to repopulate the central corneal epithelium (**Figure S7B**). In accordance with this, cluster 4 also shows high expression of *CD44* (data not shown), which is also found in migratory epithelia (F X Yu, 1998).

Cluster 9 showed an interesting transcriptional profile, exhibiting high expression of putative LSCs markers including *CXCL14, CEBPD, TP63, S100A2* and the more recently discovered marker, *TXNIP* (Kaplan et al., 2019) (**Table S1**), defining this cluster as LSCs. We selected cluster 9 to be at the start of the tree position in our pseudotime analysis, and showed that the cells gave rise to the TAs and differentiated corneal and limbal epithelial cell clusters in accordance with its LSCs definition above (**Figure S8**).

#### Identification of stromal cell populations

Fibulin 1 (*FBLN1*), which forms part of the extracellular matrix (ECM) that regulates the LSCs niche (Wang et al., 2020), was highly expressed in clusters 8 and 16 (**Figure 1**). IHC analysis, showed clear *FBLN1* expression under the limbal crypts only; however, this was not localised to the keratocytes marked by *CD34* expression (**Figure S9A**), hence, we defined cluster 8 as limbal fibroblasts. Cluster 16 displayed high expression of collagen 1 A1 (*COL1A1*) and A2 (*COL1A2*) genes, which are known to be expressed in the corneal stroma (Nakayasu et al., 1986), but more importantly this cluster also showed high expression of collagen 3A1 (*COL3A1*) (**Table S1**), whose expression is found in limbal stroma (Wang et al., 2020). A differential gene expression analysis between clusters 8 and 16, identified Osteoglycin (*OGN*) to be predominantly expressed in cluster 16 (**Figure S9B, C**). IHC analysis (**Figure S9D**) showed a clear and distinct expression of OGN in the limbal, but not central cornea, hence we defined cluster 16 as limbal stromal keratocytes.

High expression of Keratocan (*KERA*), encoding the keratan sulfate proteoglycan that is involved in corneal transparency (Kao and Liu, 2002), was found in cluster 3 and 12 (**Figure 1B, S1**). Differential gene expression analysis revealed Matrix Metallopeptidase 3 (*MMP3*) expression in cluster 3, but not 12 (**Figure S10A**). IHC localised the MMP3 immunopositive cells to the stroma of peripheral and central cornea (**Figure S10B**). All the MMP3-immunostained cells also showed expression of CD105 (**Figure S10C, C’**), a marker of CSSCs (Hashmani et al., 2013), leading us to define cluster 3 as CSSCs. Since Keratocan and Lumican (*LUM*), were amongst the top ten expressed genes in cluster 12, we performed IHC for Lumican, showing a clear “stacked arrangement” typical of keratocytes, which secrete the stroma extracellular matrix (**Figure S10D**). In view of these data we assigned cluster 12 as central stroma keratocytes.

#### Identification of immune cells

Clusters 15 and 17 were distinguished by the high expression of chemokine ligands *CCL3* in cluster 15 and *CCL5* in cluster 17 and were defined as immune cells I and II respectively (**Figure 1, S1**). Nonetheless both populations seem to have a mixed immune cell phenotype; hence, to get better insights into cell fate of these two clusters, further subclustering was performed (**Figure S11A**), revealing the presence of monocyte derived macrophages and dendritic cells, two subtypes of CD8 T cells and two subtypes of macrophages (**Table S2**), consistent with innate cell immune profiling in cornea and conjunctiva (Palomar et al., 2019).

#### Identification of endothelial cell populations

High expression of Transgelin (*TAGLN*, also known as *SM22α*; **Figure 1B**) and Actin Alpha 2 (*ACTA2*; **Figure S1**) were found in cluster 13. Both markers are associated with endothelial to mesenchymal transition, which occurs during *ex vivo* expansion of corneal endothelial cells (Roy et al., 2015), resulting in acquisition of a fibroblast cell phenotype (Roy et al., 2015). Damage to the endothelium in rabbits results in infiltration of leukocytes and loss of endothelial cells, which triggers morphological changes of surrounding endothelial cells and decreased collagen IV synthesis coupled to increased synthesis of collagens I and III (Kay et al., 1985). High levels of collagen I and III gene expression was observed, suggesting that cluster 13 represents modulated corneal endothelial cells, which have acquired a fibroblast phenotype; hence this population was assigned as fibroblastic corneal endothelial cells (FCECs). IHC indicated a small number of the FCECs to be present in the stroma of limbal and peripheral cornea (**Figure S12A**).

Clusters 11, 18 and 20 displayed high expression of the atypical chemokine receptor (*ACKR1*, also known as DARC, **Figure 1B**), whose expression is found on the endothelial cells of capillary and post-capillary venules (Thiriot et al., 2017). Differential gene expression analysis (**Figure S12B**) identified *CDH19, CCL21+LYVE1* and *POSTN* to be predominantly expressed in clusters 20, 18 and 11 respectively. IHC revealed the presence of CDH19 immunopositive cells in the corneal endothelium (**Figure S12C**) and POSTN immunopositive cells in the blood vessels in the limbus (**Figure S12D**), leading us to define clusters 20 and 11 as corneal endothelium and blood vessels respectively. Cluster 18 was defined as lymphatic vessels on the basis of CCL21 and LYVE1 expression. IHC indicated CCL21 immunopositive cells to be located in the limbal and conjunctival region (**Figure S12E**).

#### Identification of red blood cells and melanocytes

Cluster 14 showed high expression Hemoglobin Subunit Alpha 1 (*HBA1*) and other genes present in red blood cells (**Figure 1B, S1**): these were located under the conjunctival epithelium (**Figure S13A**) and the limbal crypts (**Figure S13B**). Cluster 19 showed a high expression of pigmented cell markers including *TYRP1, PMEL, MLANA, MITF* and *TYR* (**Figure 1B, S1**), which led us to define this cluster as melanocytes. IHC revealed the presence of MLANA and MITF double immunopositive cells under the limbal crypts (**Figure S13C, C’**).

In summary, our scRNA-Seq data combined with IHC analysis revealed the presence of stem, progenitor and differentiated cells in all the three layers of cornea as well as the accessory cells that are located in the limbal niche.

### scRNA-Seq analysis reveals novel markers for qLSCs and TA cells

The scRNA-Seq analysis revealed two interesting populations in the limbal epithelium, namely LSCs (cluster 9), and TA cells (cluster 4). In order to identify LSCs specific markers, we investigated which of the 119 highly expressed markers of cluster 9 (**Table S1**) were absent or expressed at low levels in the other clusters. Five markers namely *GPHA2, CASP14, MMP10, MMP1* and *AC093496.1* (Lnc-XPC-2) were highly and predominantly expressed in cluster 9 (**Figure 2A**). IHC showed distinct *GPHA2* expression all around the limbal crypt, which in some areas (bottom and top of crypt), but not all, overlapped with *KRT15* (**Figure 2B**). *GPHA2* and *MMP10* overlapped extensively throughout the limbal crypt (**Figure 2C**); however we did not observe any overlap in expression of these two markers with Ki67 (**Figure 2D**). For this reason, we will refer to this cluster as quiescent LSCs (qLSCs). MMP1 expression was also present in the limbal crypt marked by the GPHA2 and ΔNp63 expression and similarly to other markers above did not co-localise with Ki67 expression (**Figure 2E-G**). Biomart bioconductor R package was used to annotate differentially expressed genes in cluster 9 with GO terms. The glycoprotein hormone subunit alpha 2 (*GPHA2*) was annotated by GO to be located in the “cell surface” and for this reason was selected amongst the five qLSCs markers for further investigation detailed in the rest of this section. Using a similar approach, we identified *TFPI2* (tissue factor pathway inhibitor 2) to be highly and predominantly expressed (Yeh et al., 2003) in the TA cells (cluster 4) but not the other corneal epithelial cells (**Figure S14A**). IHC analysis showed a large degree of overlap between TFPI2 and ΔNp63 expression (**Figure S14B**).

**Figure 2.**
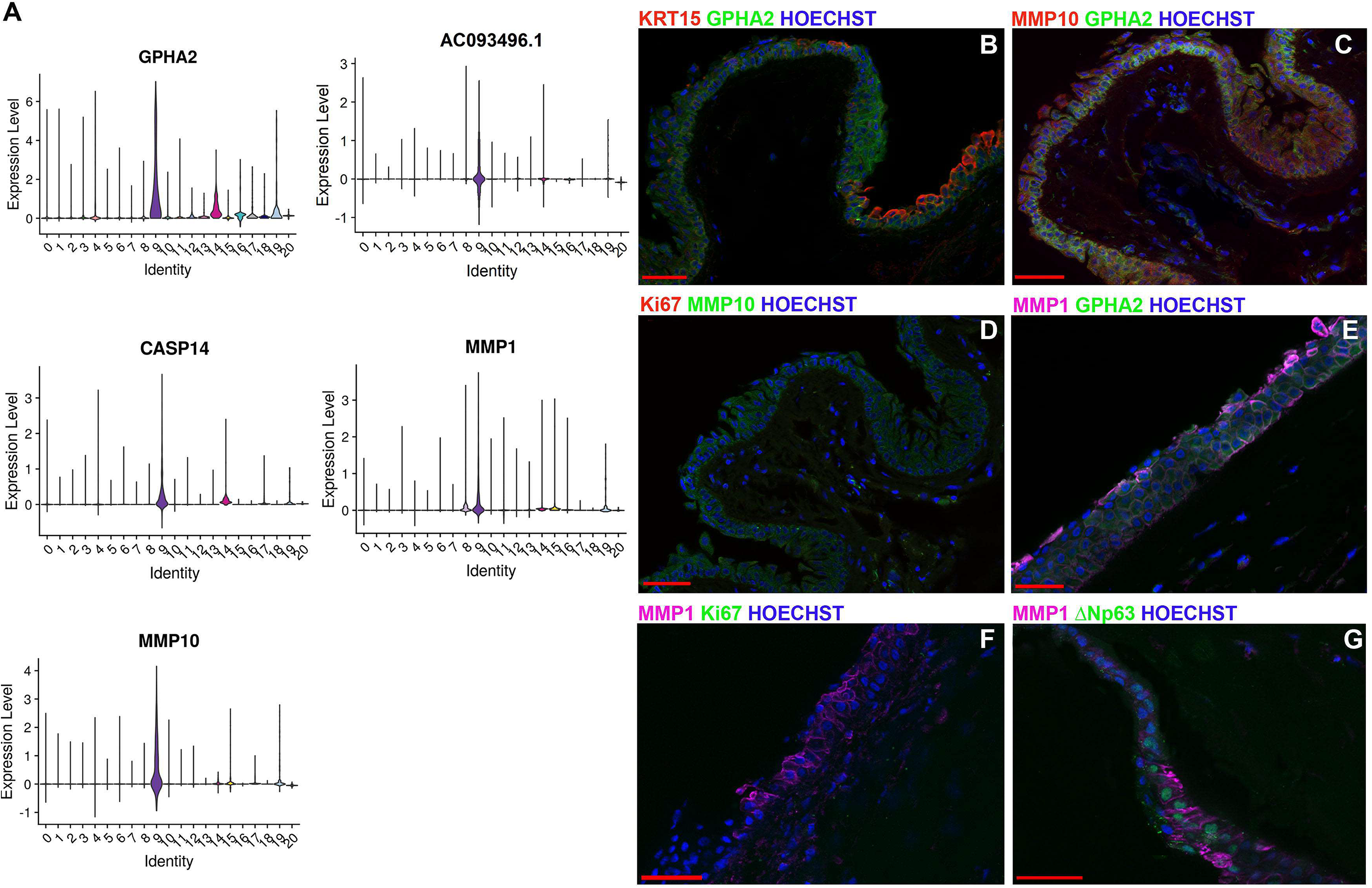
Novel markers for qLSCs. **A**) Violin plots showing the expression of five novel markers (*GPHA2, CASP14, MMP1, MMP10 and AC093496.1*), which are highly and predominantly expressed in qLSCs (cluster 9); **B-G**) IHC analysis showing the expression of GPHA2 and partial overlap with KRT15 (**B**), the overlap in expression between MMP10 and GPHA2 (**C**), lack of co-localisation between MMP10 and Ki67 (**D**), overlap between MMP1 and GPHA2 (**E**) lack of co-localisation between MMP1 and Ki67 (**F**) and overlap between MMP1 and ΔNp63 expression (**G**) in the limbal crypts. Nuclear staining indicated by Hoechst in blue colour. Scale bars: 50 μm.

Subsequently, we investigated the expression of GPHA2 and TFPI2 on *ex vivo* expanded limbal epithelial cells (LECs). GPHA2 was present in a few cells clustered together in the middle of colonies marked by KRT15 expression (**Figure 3A**). All the GPHA2 cells were Ki67^+^; however not all Ki67^+^ cells were GPHA2^+^ (**Figure 3C**). The same was also true for costaining with ΔNp63, which was observed throughout the colonies with very few GPHA2+ΔNp63+ present (**Figure 3B**) in a cluster like pattern. In accordance, flow cytometric analysis indicated 0.2 – 0.7 % of cultured LECs to display cell surface expression of GPHA2. TFPI2 was expressed throughout LEC colonies in accordance with its high putative expression in the TA cells (**Figure 3D**). In accordance with GPHA2’s expression in qLSCs, a significant reduction in expression was noticed upon air-liquid interface induced differentiation of LECs (**Figure 3E**). Flow activated cell sorting of GPHA2+ and GPHA2-cells followed by qRT-PCR analysis indicated a significantly higher expression of *GPHA2, MMP10* and *CK15* in the GPHA2+ cells, in accordance with enrichments of qLSCs on the basis of GPHA2 cell surface expression (**Figure 3F**).

**Figure 3.**
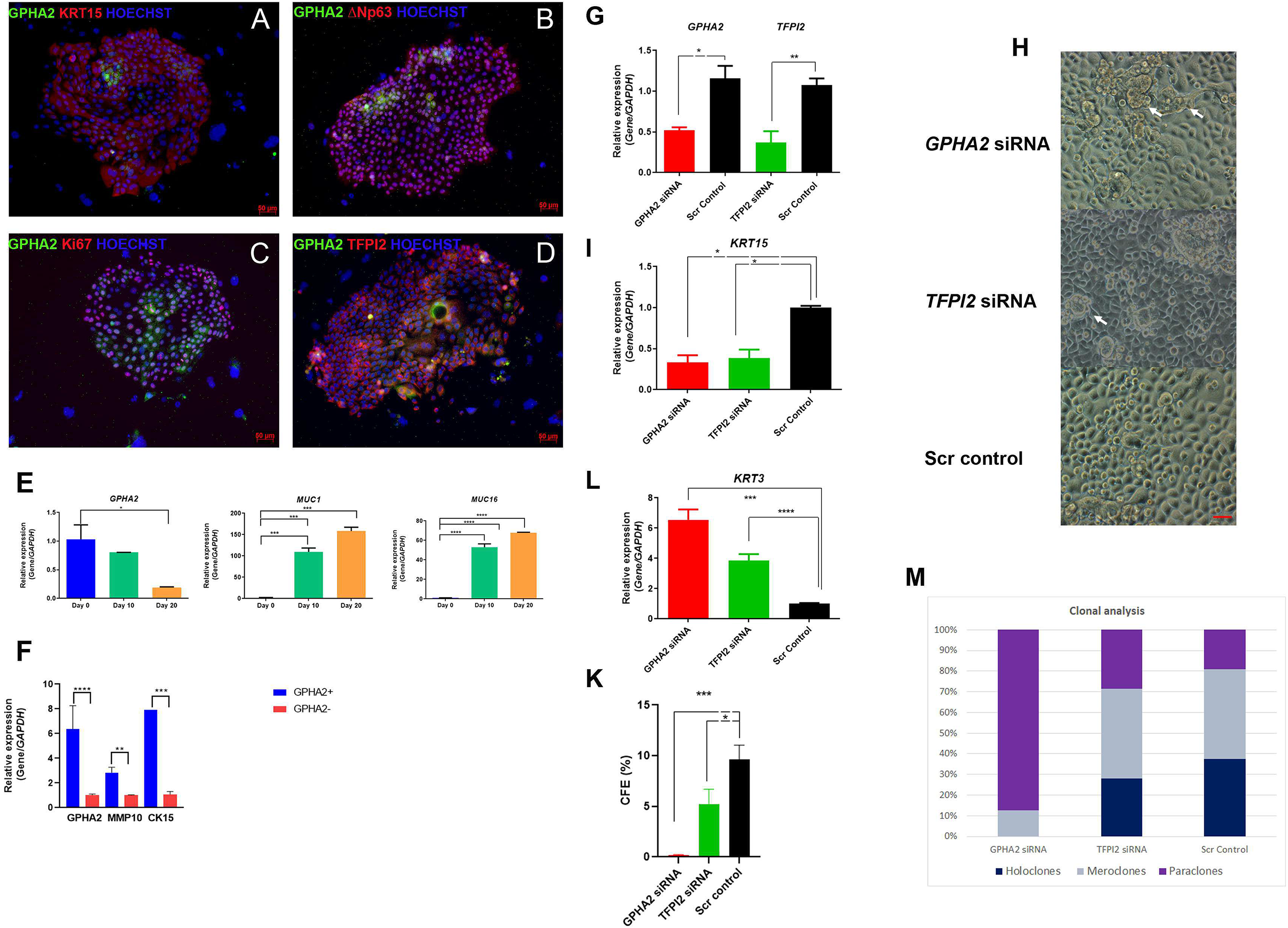
Downregulation of *GPHA2* and *TFPI2* results in loss of clonogenecity and onset of differentiation. **A-D**) Expression of GPHA2 and overlap with KRT15, ΔNp63, Ki67 and TFPI2 respectively in *ex vivo* expanded LECs, passage 1; **E**) Downregulation of *GPHA2* expression during air liquid interface culture of LECs, which was validated by the significant upregulation of limbal epithelial superficial markers *MUC1* and *MUC16;* **F**) Flow activated cell sorting combined with qRT-PCR analysis indicates enrichment of qLSCs in the GPHA2+ enriched fraction; **G**) Knockdown of *GPHA2* and *TFPI2* expression in *ex vivo* LECs using RNAi; **H**) Representative microphotographs showing the presence of larger and more differentiated like cells in the *GPHA2* and *TFPI2* siRNA treated samples, indicated by the white arrows. Scale bars 10 μm; **I, L**) The knockdown of *GPHA2* and *TFPI2* results in a significant decrease in *KRT15* (**I**) and increase in *KRT3* expression (**L**): **K**) CFE is significantly reduced upon *GPHA2* and *TFPI2* knockdown; **M**) *GPHA2* knockdown results in total lack of holoclones, a significant reduction in meroclones and increase in paraclones, whilst *TFPI2* knockdown results in a reduction in holoclones and an increase in paraclones. E-K: Data presented as mean+/− SEM, n=3. Statistical significance was assessed using oneway Anova with Dunnet Multiple Comparison Tests, * p< .05; ** p < .001, *** p < .001, **** p < .0001.

RNAi was carried out to assess the role of GPHA2 and TFPI2 in LSC clonogenecity and cell fate determination (**Figure 3G**). Morphological and qPCR analysis showed the presence of large elongated cells in the *GPHA2* and *TFPI2* siRNA groups, indicative of differentiation onset, corroborated by a significant decrease in expression of *KRT15* and the increase in expression of the differentiation maker, *KRT3* (**Figure 3H, I, L**). Colony forming efficiency (CFE) assays showed a huge reduction in the *GPHA2* siRNA group and a significant decrease in the *TFPI2* siRNA group (**Figure 3K**). Clonal analysis showed a complete lack of holoclones upon *GPHA2* knockdown (**Figure 3M**), with the majority of the colonies being completely differentiated with a typical paraclone appearance. A small but significant reduction in holoclones was present in the *TFPI2* siRNA group and this was compensated by an increase in paraclones (**Figure 3M**).

Together these data indicate an important role for GPHA2 and TFPI2 in qLSCs and TA selfrenewal and differentiation.

### Combined scRNA-Seq and ATAC-Seq analysis reveals key transcriptional networks in qLSCs and TA cells and close interactions with immune cells

The same human adult cornea and conjunctiva samples used for the scRNA-Seq analysis were also subjected to scATAC-Seq analysis. Approximately 10,000 cells were captured using the 10 X Chromium Single Cell ATAC Library & Gel Bead Kit Genomics (version 1). Following QC, 10,625 cells were obtained after data integration. All the predicted RNA-Seq clusters were also found in ATAC-Seq analysis with the exception of cluster 20, which may be due to the very low cell numbers present in this cluster (**Figure S15**). Interestingly, the qLSCs and TA cell clusters were distinct in the UMAP plot from the other epithelial clusters. To analyse these in more detail, 7618 epithelial cells were selected (**Figure 4A**) and differential accessibility analysis of the qLSCs (cluster 9) or TA cells (cluster 4) versus the rest of the corneal and conjunctival cell clusters was performed. This analysis identified differentially accessible (DA) peaks overlapping with promoters (−1000bp, +100bp) and distal or intergenic regions of any transcription start site (**Table S3, S4**). Amongst the top accessible promoters in qLSCs were those of *KRT14* and *CASP14* (**Figure 4B, C**), both highly expressed in this cluster (**Figure S5B, 2A**). Similarly, *TFPI2* was amongst the top accessible promoters of cluster 4, (**Figure 3D, 4D**). Amongst the less accessible promoters in both qLSCs and TA clusters 9 and 4, we identified *MUC22* (**Figure 4E**), which is highly expressed in the corneal and limbal superficial epithelial clusters (**Figure 1A**), but not in qLSCs and TA cells.

**Figure 4:**
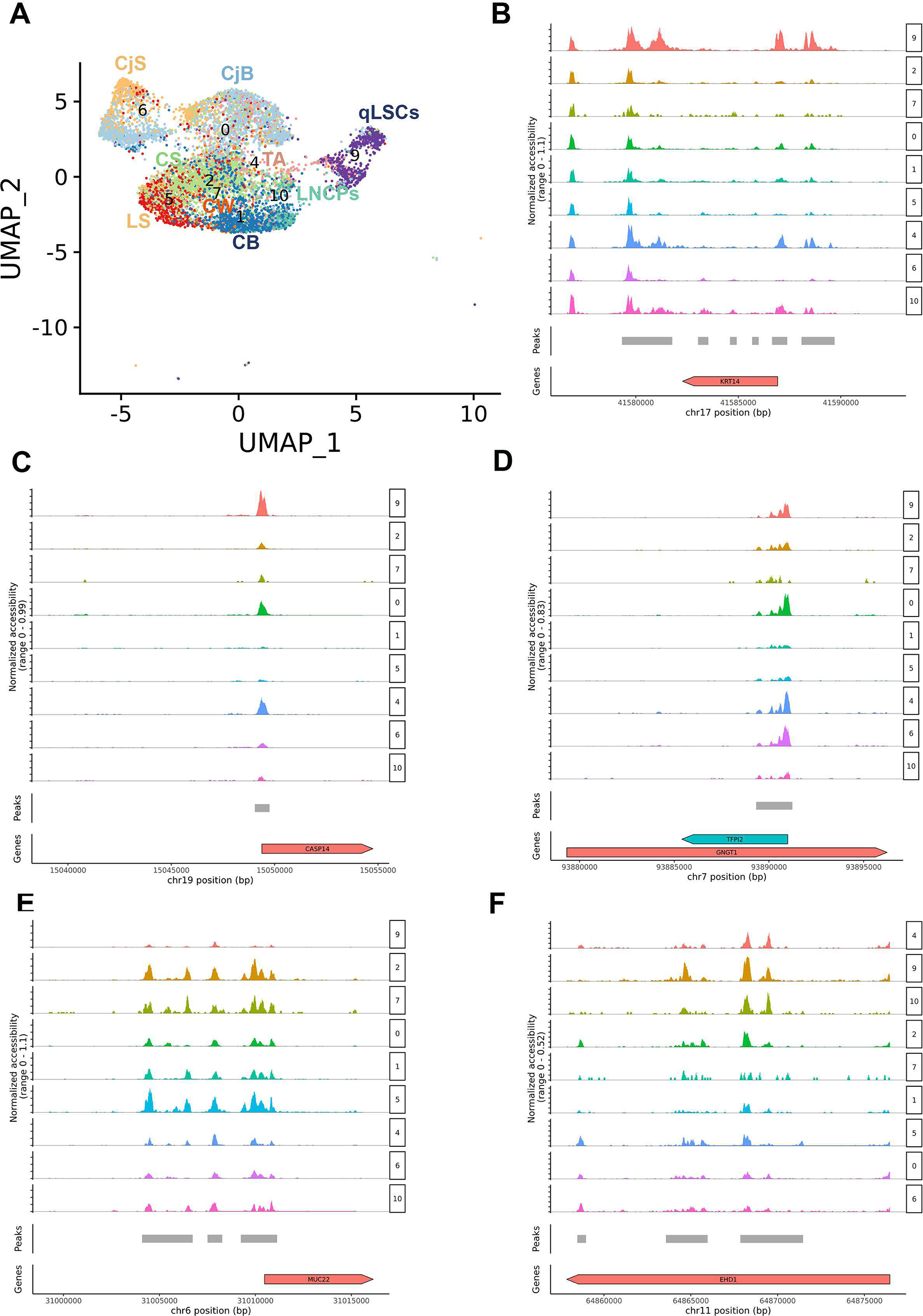
scATAC-Seq of adult human cornea and conjuctiva. (see also Figure S15, Tables S3, S4). **A**) UMAP of adult human cornea and conjunctiva epithelial clusters and limbal neural crest progenitors; **B-F**) Schematic single cell chromatin accessibility of *KRT14* (**B**), *CASP14* (**C**), *TFPI2* (**D**), *MUC22* (**E**) and *EHD1* (containing a distal enhancer for *GPHA2*) (**F**) in the human cornea and conjunctiva epithelial clusters and LNCPs.

The JEME database (Cao et al., 2017) was used to link the differentially accessible peaks to the enhancer regions for qLSCs (cluster 9) and TA cells (cluster 4). The enhancer of the qLSCs unique marker, *GPHA2*, was identified as being more accessible in cluster 9 (**Figure 4F**), in accordance with its function in maintaining qLSCs self-renewal. Transcription factor (TF) binding motifs enrichment was performed for each cluster using Signac (**Table S3, S4**). It is of interest to note that binding factor motifs for the putative LSCs and progenitor markers, such as *TP63* and *CEBPD*, were enriched in both qLSCs and TA cell clusters (**Table S3, S4**).

To identify TF networks governing the qLSCs and TA cell clusters the upstream regulator tool in IPA was used, combined with overlay analysis of DA peaks, enhancers and TF binding motifs. This analysis revealed enhanced activation of 23 upstream regulators in qLSCs compared to the rest of epithelial clusters (**Table S5**). Fourteen out of the 23 upstream regulators represent pro-inflammatory cytokines (TNF, IL1β, IL6, IL17A, IFN□ and OSM), pro-inflammatory cell surface receptors (TREM1), inducers of proinflammatory cytokine expression (AP1) and regulators of inflammatory response (NFkβ, RELA, CSF2, PI3K, ERK1/2, and F2). Importantly, 6 of these regulators (*TNF, IL1B, IFN□, OSM, TREM1, CSF2*) show the highest expression in immune cells 1 and/or 2 clusters (**Figure S16**), whilst a further regulator (IL6) is predominantly expressed in immune cells and limbal fibroblasts (data not shown). Importantly, CellPhoneDB (Efremova et al., 2020), revealed multiple significant interactions between cluster 9 and immune cells (clusters 15 and 17, **Table S6**). Together these findings suggest an important role for pro-inflammatory cytokines produced by the immune cells in the regulation of qLSCs quiescence and activation. This hypothesis is corroborated by published bulk RNA-Seq data and supplementation of cultures with some of these pro-inflammatory markers impacting directly on expression of putative LSC markers and their colony-forming efficiency and size, which serve as independent validation benchmarks for our TF network analysis. (Veréb et al., 2013, Yang et al., 2019, (Zhu et al., 2020), Notara et al., 2010). This is fully supported by *in vivo* mouse data, which show that the absence or inhibition of immune cells, results in loss of LSCs quiescence and delayed would healing (Ruby Shalom-Feuerstein, personal communication) as well as the clinical phenotype of immune-mediated genetic ocular surface diseases (e.g. Steven-Johnson syndrome, allergic chronic limbitis or ocular cicatricial pemphigoid) and chronic ocular surface inflammation, where excess inflammation can lead to LSCD.

Amongst the upstream regulators, we also found a number of growth factors, namely EGF, BDNF and FGF2 (**Table S5**). Addition of EGF has become a standard media requisite for the expansion of LECs, with EGF addition stimulating proliferation (Trosan et al., 2012), colony growth under serum free conditions (Meyer-Blazejewska et al., 2010) and inhibiting the expression of differentiation markers (Wilson et al., 1994). EGFR is present in qLSCs (Zieske et al., 1993) (**Table S6**) and its inhibition has been shown to affect epithelial cell proliferation during corneal wound healing (Nakamura et al., 2001). Although a direct role for FGF2 in limbal stem cell expansion has not been proven, FGFR2 is essential for corneal epithelial cell proliferation and differentiation (Zhang et al., 2015). Similarly, the nerve growth factor BDNF, may influence qLSCs, via promotion of corneal innervation (Ueno et al., 2012). One of the BDNF receptors, the low affinity nerve growth factor receptor, p75, is differentially expressed in LSCs and downregulated during differentiation (Kolli et al., 2019). Impairment of trigeminal innervation, which provides trophic support to the cornea results in neurotrophic keratitis, a degenerative disease characterized by corneal sensitivity reduction, spontaneous epithelium breakdown, and impairment of corneal healing (Sacchetti and Lambiase, 2014). The presence of BDNF as an upstream regulator in qLSCs may therefore underline the close interaction between qLSCs and corneal nerves, contributing to the maintenance of corneal epithelial surface integrity and consequently ocular surface homeostasis.

It is interesting to note that pro and anti-angiogenic factors including Coagulation Factor 2 (Thrombin), VEGF and PDGF BB were found as upstream regulators in qLSCs. While VEGF and PDGF BB are pro-angiogenic factors (Cursiefen et al., 2000), thrombin itself can activate thrombospondins 1 and 2 (TSP-1 and TSP-2), which are expressed in the cornea and contribute to its avascularity (Cursiefen et al., 2004). Collectively, these data suggest a balanced action of pro- and anti-angiogenic regulators in qLSCs to maintain an avascular state in the corneal epithelium.

Most of these upstream regulators involved in inflammatory signalling, angiogenic and growth factor based are also present in the TA cells (cluster 4). However, new additional regulators including the putative LSCs marker TP63, transcriptional regulators TP53, GLI1, ECM components (e.g. fibronectin), hormones (e.g. insulin) etc. were found amongst the upstream regulators of gene expression in cluster 4. Given the large number of upstream regulators, it is impossible to analyse and validate each one in detail; however, we anticipate that provision of these regulators for qLSCs and TA cells would provide a useful platform for further studies.

### LECs expansion *ex vivo* and limbal dysplasia *in vivo* result in downregulation of qLSCs and acquisition of proliferative limbal progenitor markers

*Ex vivo* expansion of LECs is widely used for treatment of patients with LSCD either by single cell disassociation of limbal rings and plating on mitotically inactivated 3T3 fibroblasts or explant outgrowth on a substrate such as human amniotic membrane (HAM) (Baylis et al., 2011). Although these techniques are widely used clinically, it is not known whether the *ex vivo* expanded LECs resemble the qLSCs or TAs *in vivo* or if either technique is superior. To address these questions, we obtained one cadaveric limbal ring, which we analysed by scRNA-Seq before and after *ex vivo* expansion on 3T3 feeders and HAM. 14,897 cells were obtained after filtering and QC steps. To facilitate cell cluster annotations, this dataset was integrated with the four adult human cornea/conjunctival datasets using Seurat. This method is designed to overcome batch effects and identify shared cell identities across different experiments, whilst preserving the identity of cells that are unique to a specific cluster. We identified three additional clusters in the *ex vivo* expanded LECs (**Figure 5A** and **Table S7),** all of which were characterised by a high percentage of cells in S and/or G2/M phase of the cell cycle (**Figure 5B**) and high expression of *Ki67* and *PCNA*, indicative of their proliferative nature (**Table S7**).

**Figure 5:**
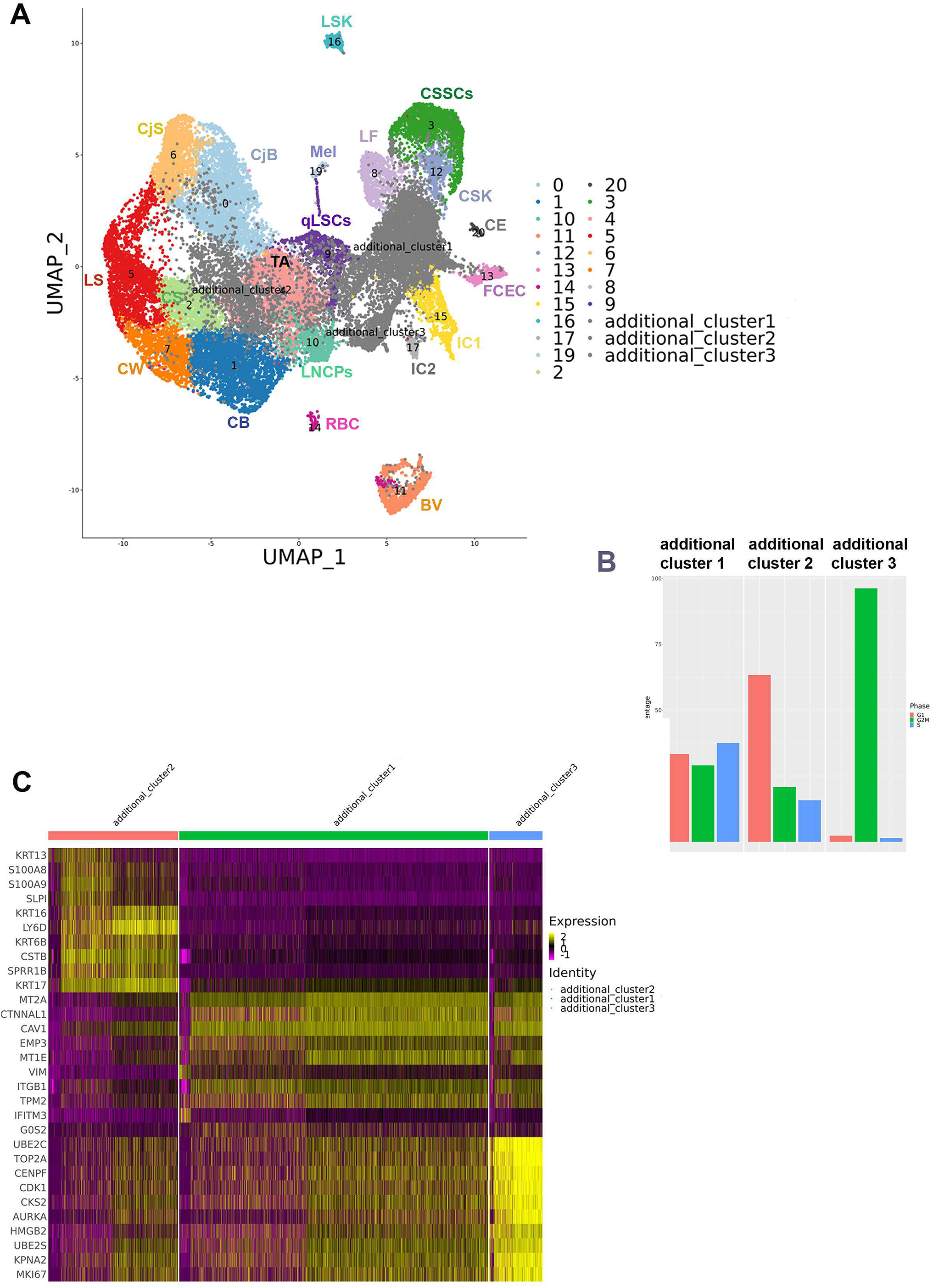
scRNA-Seq of *ex vivo* expanded human LECs and comparison to adult human cornea and conjunctiva. (see also Figure S17 and Table S7**. A**) UMAP showing the presence of three additional cell clusters found in the *ex vivo* expanded LECs. All cluster annotations are the same as in Figure 1A; **B**) Cell cycle distribution of additional clusters 1-3; **C**) Comparative heatmap showing differentially expressed genes between the three additional clusters found in the *ex vivo* expanded LECs.

Differential gene expression analysis identified several markers associated with limbal basal epithelium, including *ITGA6* (Thomas et al., 2007)*, ITGB1* and *VIM* (Schlötzer-Schrehardt and Kruse, 2005) in the additional cluster 1, defining it as cultured basal limbal epithelium (**Figure 5C**). Markers associated with basal (*KRT17*) and conjunctival epithelium (*KRT13, KRT7*) were most highly expressed in the additional cluster 2 (**Table S7**), hence, this cluster was annotated as cultured basal conjunctival epithelium. The presence of such a cell cluster is not surprising, as limbal ring dissections can often include a small rim of surrounding conjunctiva. The additional cluster 3 was characterised by high expression of markers of mitosis in addition to epithelial progenitor markers *KRT14* and *KRT15* and was therefore annotated as mitotic epithelial progenitors. A correlation analysis was performed by taking the average gene expression of each cluster for the top 2000 highly variable genes used in the clustering analysis. This analysis indicated the additional cluster 1 to be more similar to the qLCSs cluster 9 (correlation coefficient=0.91), whilst the additional cluster 2 was similar to both TA cluster 4 and qLSCs cluster 9 (correlation coefficient 0.93 and 0.92 respectively).

Culture on 3T3 feeders and HAM generated similar percentages of basal limbal, conjunctival epithelial cells and mitotic progenitors (Figure **S17A)**. Differential gene expression analysis was used to compare expanded cultured basal limbal cells (additional cluster 1) to qLSCs *in vivo* (**Figure S17B**), revealing a significant decrease in the expression of qLSCs markers (**Table S8**) including *GPHA2, MMP10, CASP14, TXNIP* and *CEBPD*, corroborating the presence of very few GPHA2 cells in *ex vivo* cultured LECs (**Figure 3A**). Similarly, significant downregulation of *KRT13, KRT19* and *KRT15* was observed when basal conjunctival cells *in vivo* (cluster 0) were compared to the *ex vivo* expanded basal conjunctival epithelial cells (**Table S8**). In addition a significant upregulation of markers associated with highly proliferative limbal progenitor cells (e.g. *S100A2, S100A10*) (Li et al., 2011) was observed when the additional cluster 1 was compared to the qLSCs.

To investigate if a similar phenomenon occurs during dysregulation of LSCs growth *in vivo*, we focused on scRNA sequencing of an adult cornea with a visible limbal growth/dysplasia protruding from the limbus towards the nasal conjunctiva (**Figure 6A**). The scRNA-Seq subset obtained from the cornea with limbal dysplasia was integrated with the adult cornea and conjunctiva and cultured LEC subsets (**Figure 6B**), resulting in the identification of six additional clusters (**Table S9**), five of which were present in both the cornea with limbal dysplasia and cultured LSCs (**Figure S18**).

**Figure 6:**
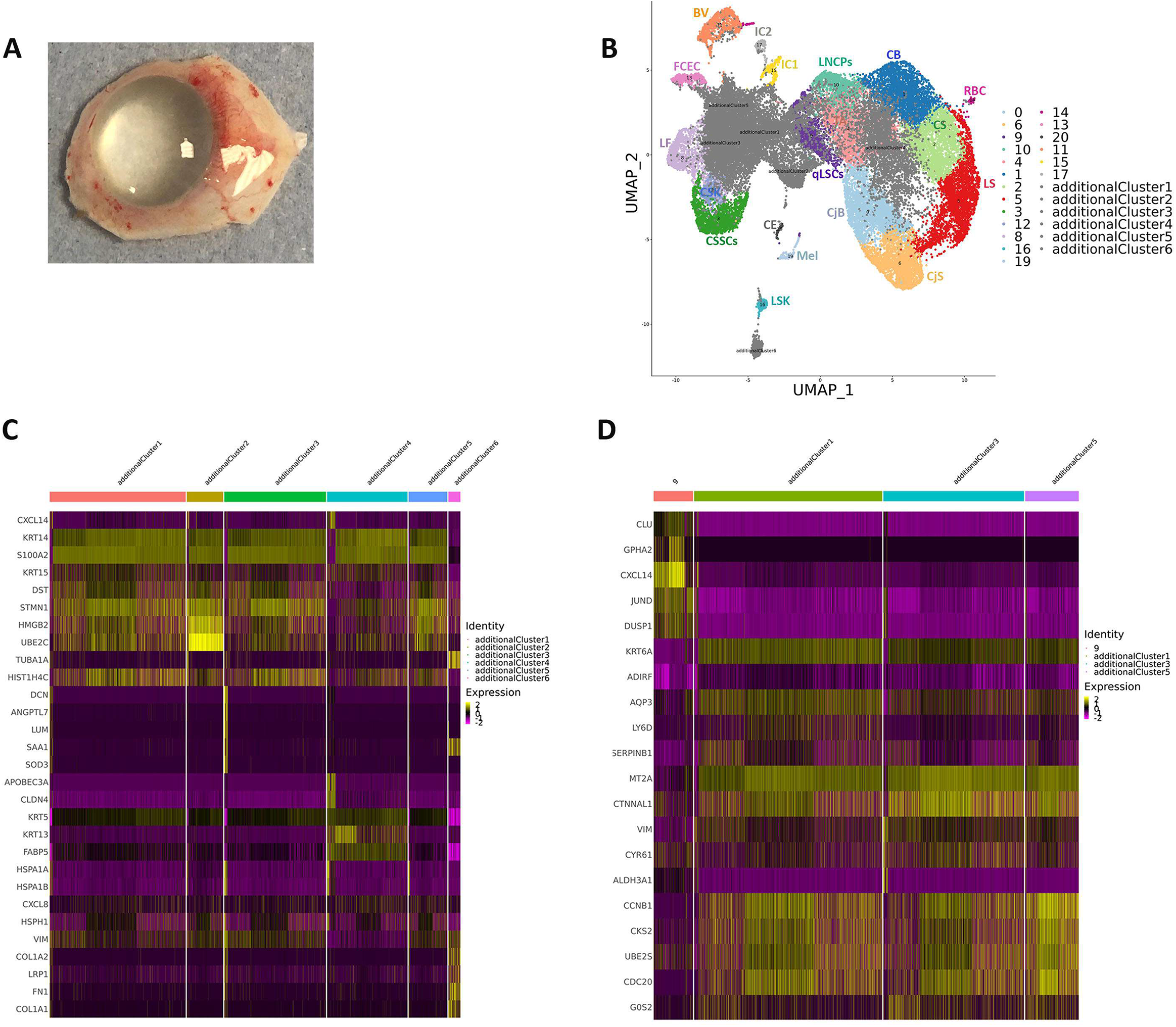
Expansion of LECs *in vivo* leads to loss of qLSCs and acquisition of markers associated with proliferative limbal epithelial progenitors. (see also Figures S18, S19 and Table S9)**. A**) Representative photo showing a human cornea with a limbal dysplasia. All cluster annotations are the same as in Figure 1A; **B**) UMAP showing the presence of six additional clusters in the cornea with limbal dysplasia; **C**) Comparative heatmap showing the differentially expressed genes between the six additional clusters found in the cornea with dysplasia; **D**) Comparative heatmap showing differentially expressed genes between qLSCs (cluster 9) and the proliferative basal limbal epithelium I-III corresponding to additional clusters 1, 3, 5 in the cornea with limbal dysplasia.

The additional cluster 6, which is specific to the cornea sample with limbal dysplasia, displayed high expression of *SAA1*, a marker of pancreatic ductal adenocarcinoma tumor stroma primarily composed of cancer-associated fibroblasts (Djurec et al., 2018) (**Table S9**). SAA1 is also expressed in the corneal fibroblasts and corneal keratocytes and shown to be upregulated in inflammation mediated neovascularisation (Liu et al., 2011). This cluster also displayed high expression of other fibroblast and keratocytes markers including *VIM, FN1, COLA1A2, TIMP2* (**Figure 6C** and **Figure S19C**) and for this reason was defined as activated fibroblastic stroma cluster.

The proliferation markers, *PCNA* and Ki67 were highly expressed in additional clusters 1-5, indicating their proliferative nature (**Figure S19A**). Cell Cycle analysis corroborated these findings (**Figure S19B**) showing a considerable percentage of cells in the S and/or G2/M phases of the cell cycle. Differential gene expression analysis indicated high expression of proliferative limbal epithelial progenitor markers (*KRT14, KRT15, S100A2* etc.) in the additional clusters 1-5 (**Figure S19C**). Additional cluster 4 showed high *KRT13* expression, a typical marker of conjunctival epithelium (**Figure 6C**) and thus was defined as proliferative conjunctival epithelium. Cluster 2 contained more than 95% of cells in mitosis and was defined as mitotic epithelial progenitors. The additional clusters 1,3 and 5 were defined as proliferative basal limbal epithelium I-III in view of their cell cycle characteristics, high expression of basal limbal markers (*CXCL14* etc) and lack or minimal of expression of the conjunctival marker, *KRT13* (**Figure 6C**). Similarly to *ex vivo* expanded LECs, the additional clusters 1, 3 and 5 showed a downregulation of markers associated with qLSCs including *GPHA2, MMP10, CASP14, TXNIP* (**Figure 6D**), indicating that limbal dysplasia *in vivo* may also result in downregulation of typical qLSCs and acquisition of proliferative limbal progenitor markers.

### scRNA-Seq of keratoconus corneas reveals activation of collagenase in the corneal stroma and a reduced pool of TA cells in the limbal epithelium

To validate the applications of single cell sequencing as a platform for gaining quick insights into disease pathology we focused on keratoconus, an asymmetric, progressive disease in which the cornea becomes conical in shape. The aetiology of keratoconus is not fully understood although current knowledge postulates that this is a final common pathway for several diseases, which are underlined by genetic predisposition triggered by environmental factors (Mas Tur et al., 2017). To gain insights into disease pathology we performed scRNA-Seq of central cornea samples obtained from two affected human subjects with (18 and 43 year old males) at the time of corneal transplantation and one cadaveric unaffected female adult subject. After filtering and QC steps, 2,641 cells were obtained. This dataset was integrated with the adult cornea/conjunctiva dataset (**Figure 7A**), resulting in identification of a new additional cluster of central stroma keratocytes (**Table S10**), which was present in the central cornea of both unaffected subject and keratoconus patients. The percentage of cells in each cluster (**Figure 7B**), revealed a noticeable decrease in TA cells (cluster 4) and an increase in cluster 12 (corneal stroma keratocytes).

**Figure 7:**
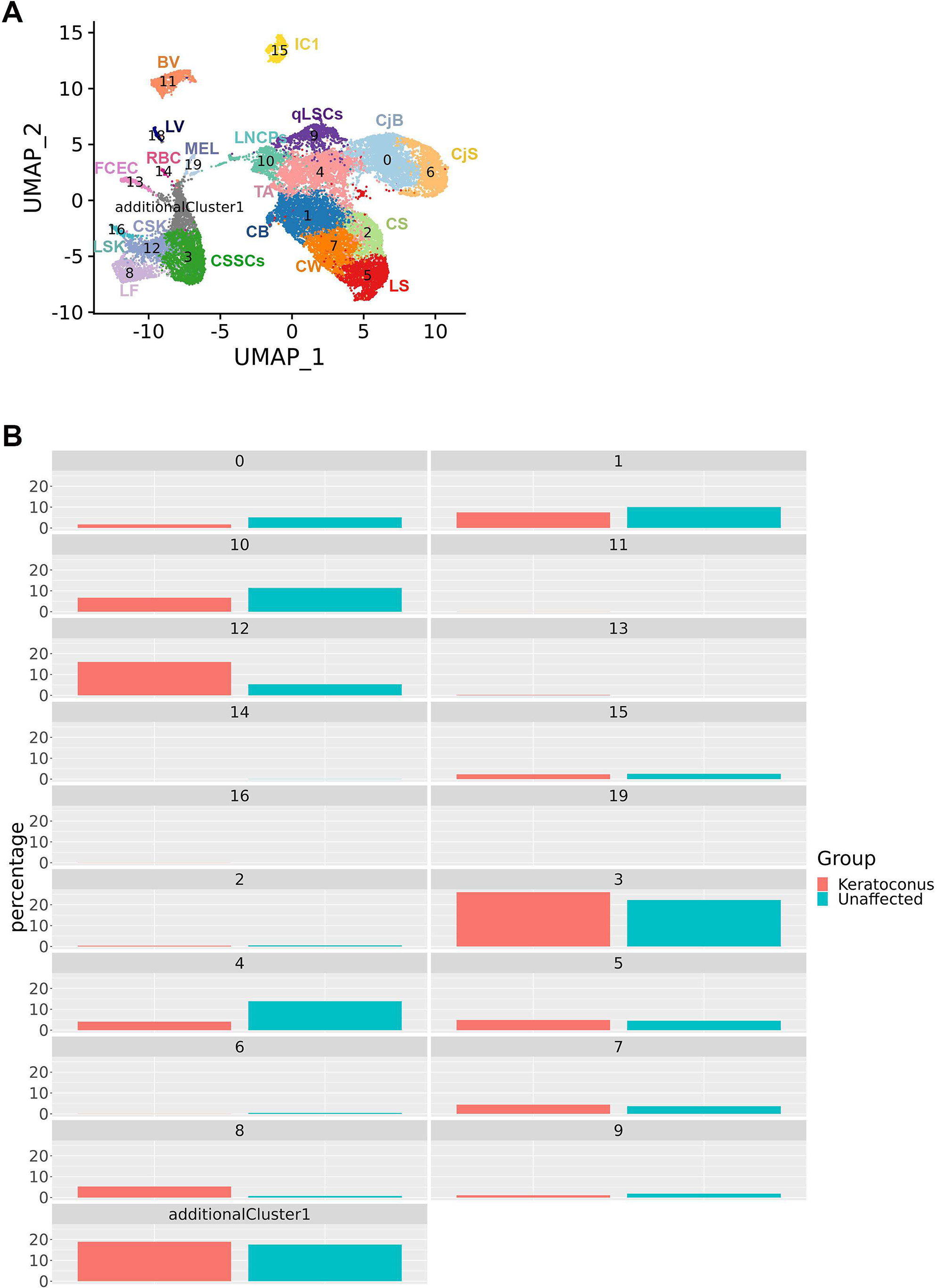
scRNA-Seq of keratoconus samples. (see also Figures S20, S21 and Tables S10-S13)**. A**) UMAP of cornea and conjunctiva samples obtained from two patients with keratoconus and one unaffected subject; **B**) Comparative analysis between the unaffected and keratoconus cornea and conjunctiva showing the percentage of cells in each cluster.

Irregular arrangement, enlargement and reduction in corneal basal epithelial cell density of keratoconus patients is known (Khaled et al., 2017). In addition, loss of corneal stromal architecture is a key feature of keratoconus affected corneas. In view of this and the changes noted in cluster 12, we performed differential gene expression analysis for this cluster (**Figure S20A,B, Table S11**), which indicated a significant downregulation of genes involved in collagen biogenesis (*COL8A1, COL6A3, COL5A2*), proteoglycans (*KERA*), genes involved in WNT signalling pathway (*WNT5A, DKK2, SFRP1*), serine protease inhibitors (*SERPINA5, SERPINE2*), mitochondrial genes (*MT-CO1,MT-ND1*), cytokeratins (*KRT5, KRT14*), cytokines (*IL32*) and members of the TGFβ (*TGFβI*) family involved in collagen-cell interaction in keratoconus stroma samples, corroborating previously published data (Chaerkady et al., 2013; Khaled et al., 2017; Mas Tur et al., 2017; Shinde et al., 2019). In contrast, we noticed a significant upregulation of genes involved in ECM degradation (*TIMP3, TIMP2*), cell death (*ANXA1*), oxidative stress defence and detoxification (*ALDH3A1*), epithelial to mesenchymal transition (*VIM, TWIST1*) and FOS and JUN oncogenic transcription factors (*FOS, FOXB, JUNB, JUND*) (**Figure S20B**).

The top canonical pathways that were most significant to keratoconus stroma were EIF2 and mTor signalling, oxidative phosphorylation, and mitochondrial dysfunction (**Table S12**). Interestingly these top pathways have been associated with keratoconus pathogenesis through proteomic studies of corneal stroma obtained from affected patients (Chaerkady et al., 2013; Shinde et al., 2019; Vallabh et al., 2017; Wojcik et al., 2013), providing a validation benchmark for our single cell studies.

Using IPA, we identified activated upstream regulators with the most significant for Keratoconus being collagenase (**Table S13**). The increase in collagenase activity, which has been biochemically confirmed in keratoconus stroma, is of great interest as it has been shown that this enzyme cleaves collagen molecules into fragments, which are processed by gelatinase (also activated in keratoconus stroma) and cathepsin for degradation, corroborating the observed decrease in collagen in the corneal stroma of keratoconus patients. Together our scRNA-Seq studies support the original hypothesis of Kao et al that the decrease in collagen and stromal thinning in keratoconus is due to an increase in collagenase activity (Kao et al., 1982); therefore keratoconus may represent a collagenolytic disease. The loss and degradation of stromal collagen may allow other cell types to repopulate the corneal stroma in keratoconus patients. Based on the significant increase in expression of *TWIST1*, a key transcription factor driving epithelial to mesenchymal transition and *VIM*, a key marker of mesenchymal cells, we suggest that the non-resident stromal populating cells may be of mesenchymal origin.

The typical presentation of keratoconus has both stromal and central epithelial thinning (Khaled et al., 2017); however the underlying cause for the epithelial thinning is not known. Given the decrease in percentage of cells observed in the TA cell cluster 4 in keratoconus patients (**Figure 7B**), we performed a differential gene expression analysis (**Figure S21, Table S11**), which indicated a significant decrease in epithelial progenitor and basal cell markers (*KRT14, KRT15, IFITM3, GJA1, TXNIP*) as well as a significant increase in expression of differentiated epithelial cell markers (*AREG, KRT3, HES1*) in keratoconus samples. Together these data suggest that in keratoconus patients, the TA cells differentiate towards wing/superficial epithelial cells, depleting the pool of migratory TA cells that are able to repopulate the central corneal epithelium. Given previous interactions between immune and TA cells described earlier and the dominance of inflammatory driven signalling pathways in TA cells in keratoconus patients (**Table S11**), it may be possible that these changes are driven by inflammatory processes, also observed in tears of keratoconus patients (McMonnies, 2015).

### scRNA-Seq of human developing cornea reveals stage-specific definitions of corneal epithelial, stromal and endothelial layers

To understand the molecular events that lead to specification of stem and progenitor cells in the epithelial, stromal and endothelial layers of the cornea, we performed scRNA-Seq analysis on seventeen human corneas dissected from 10-21 post conception week (PCW) specimens. Following filtering and QC, 89,897 cells were analysed. The expression of typical epithelial, stromal and endothelial cell markers was not detectable at the very early stages of corneal development (Davies et al., 2009; Rodrigues et al., 1987); hence, we relied on published evidence of developmental tissue markers contributing to the corneal layer development (neural crest, periocular mesenchyme, mesoderm) and the transfer label function from Seurat to project cell annotations from the ***adult cornea*** into the developmental samples.

At 10 PCW, a large cell cluster of neural crest cells was identified alongside a smaller cluster of mesodermal and proliferating progenitors (**Figure 8**), showing high expression of *IGFBP5/PITX2/FOXC2* (Gage et al., 2005)*, PITX1* and *Ki67* respectively (**Table S14**). At this stage of development, the predominant epithelial cluster is the ocular surface epithelium, characterised by high expression of several cytokeratins (17, 18, 19, 8), mucins (15, 16, 20) and Metallothionein (*MT2A, MT1X*). During 6.5-21 PCW, the ocular surface epithelium specifies into limbal, corneal and conjunctival epithelium; however, the order of these events is unknown in molecular terms (Davies et al., 2009b). Our data suggest that by 10 PCW, the first of these events has occurred with conjunctival epithelium (characterised by high expression of cytokeratin 13 and several mucins (1, 15, 16, 20)) being detected as a small cluster. Similarly to ocular surface epithelium, conjunctival epithelium expresses high levels of several other cytokeratins (4, 5, 8, 17, 18, 19); however, neither the corneal nor the conjuctival epithelium express high levels of cytokeratin 3, 12 or any of the progenitor or qLSCs markers (*CK15, CK14, TP63, GPHA2, MMP10* etc) at this stage. The stromal compartments (keratocytes) were easily identified by the high expression of keratocyte markers, *TGFBI and THBS1* as well as the expression extracellular matrix components (*KERA, LUM*, various collagen chains) secreted by the corneal stromal keratocytes (**Table S14**) (Sevel et al., 1988). The endothelial cluster was distinct from the others and expressed high levels of several genes found in developing endothelial cells including *KDR, MSX1, BMP2* and *COL4A2*.

**Figure 8:**
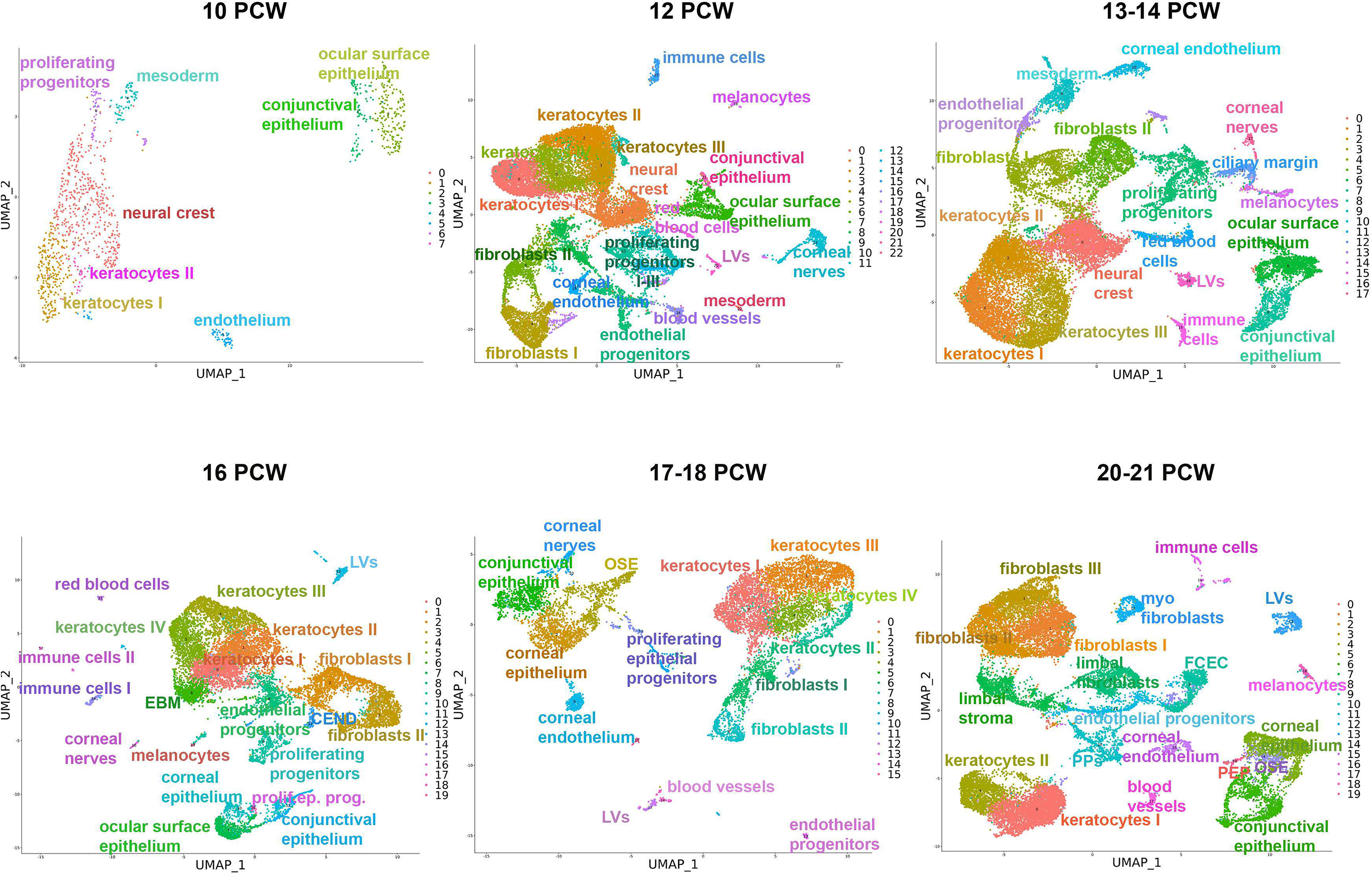
scRNA-Seq of embryonic and fetal cornea and conjunctiva from 10 to 21 PCW with cluster annotations (see also Table S14). *Abbreviations:* CEND – corneal endothelium EBM – epithelial basement membrane Ep – epithelial LV – lymphatic vessels FCEC – fibroblastic corneal endothelial cells PEP – proliferating epithelial progenitors Prolif – proliferating Prog – progenitors OSE – ocular surface epithelium

A much higher complexity was observed at 12 PCW, where the number of detected cell clusters reached 23 (**Table S14**). In addition to neural crest cell and mesodermal clusters, a higher complexity of corneal stromal populations was observed with increased cellularity (**Figure 8**) alongside increased complexity of fibroblast cell clusters. The ocular surface epithelium still forms the larger cluster compared to conjunctival epithelium, which is defined by cytokeratin 13 expression. The ocular surface epithelium displays high expression of *KRT15, KRT14, TP63, PAX6*, corroborating published evidence of p63 and CK15 immunostaining throughout the corneal epithelium at this developmental stage (Davies et al., 2009). Three clusters of proliferating cells were identified, indicative of ongoing cellular proliferation and corneal immaturity. At this stage of development, corneal endothelium and endothelial progenitor cells were present alongside two additional clusters representing the blood and lymphatic vessels (**Figure 8B**). This is also the first developmental stage in which melanocytes, red blood cells, corneal nerves and immune cells were also detected (**Table S14**). The 13-14 PCW looks very similar to 12 PCW in terms of cluster identity and the presence of accessory cells types (**Figure 8**). At this stage of development, the ocular surface epithelium continues to show high expression of *CXCL14, KRT15, KRT14, KRT19, KRT15, TP63* and secreted WNT family members (*WNT6A, WNT10A, WNT4, WNT7B, WNT2B*) (**Table S14**). Similarly to 12 PCW stage of development, the expression of qLSCs markers was not observed, indicating that limbal epithelium has not been specified yet.

At 16 PCW, the neural crest and mesodermal cell clusters are no longer visible, suggesting their migration and differentiation to corneal stromal and endothelial cell layers is largely completed (**Figure 8**). Two major differences are seen compared to 10-14 PCW. Firstly, the detection of a cell cluster (6) with high expression of Nidogen (**Table S14**), which may suggest the development of the corneal epithelial basement membrane, corroborating histological observations of a thin Bowman’s layer in 15 PCW specimens (Davies et al., 2009). Secondly, corneal epithelium (distinguished by *KRT12* expression) and a cluster of proliferating epithelial progenitors with high *KRT15, KRT17, KRT19, KRT18, KRT14* and *WNT7B, WNT4, WNT10A* and *WNT6* expression is now obvious in addition to a remaining smaller cluster of ocular surface epithelium and already defined conjunctival epithelium cluster. The proliferating epithelial progenitor cluster shows high expression of TA markers *TP63, CLDN1, CLDN4* and qLSCs marker *TXNIP*, which suggests some demarcation of “a peripheral limbal like region” harbouring the limbal stem and/or progenitor cells. This is in accordance with scanning electron microscopy studies, revealing a clear demarcation zone containing CK15 immunopositive cells between the smooth cornea and conjunctiva at 16 and 17 PCW fetal specimens (Davies et al., 2009).

The 17th and 18^th^ week of development are characterised by very similar cell clusters to 16 PCW, however, there is increased cellularity of corneal and conjunctival epithelium compared to the other developmental stages (**Figure 8**). At 20-21 PCW, the ocular surface epithelium cluster is much smaller than in previous stages, indicating that most of the commitment to the defined corneal, limbal and conjunctival epithelium has already taken place. In accordance with the detection of the proliferating epithelial progenitor cell cluster expressing TA and qLSCs markers, clusters of limbal fibroblasts and stroma were identified, suggesting the formation of a specialised limbal niche to support the self-renewal and differentiation of limbal and progenitor stem cells. These findings correlate with published histological evidence showing the presence of a loosely organised limbal stroma prior to the formation of the limbal ridge (Davies et al., 2009). At this stage of development, an increase in fibroblast type and cellularity is observed. In addition a defined cluster of myofibroblasts is now identified (**Figure 8**). The myofibroblast’s key role is the restoration of corneal integrity because of their ability to secrete extracellular matrix, contribute to wound repair and adhesion capability to the surrounding substrates. In adults, development of corneal myofibroblasts is noticed after surgery and is considered as a pathological response to injury. Their presence at the midgestation stage may suggest a “beneficial” role in response to apoptosis that cornea is thought to undergo during morphogenesis, resulting in specification of the limbal ridge (Davies et al., 2009). We were able to detect for the first time during corneal development the FCECs cluster, which is characterised by the expression of markers involved in endothelial to mesenchymal transition, a process known to occur during *ex vivo* expansion of corneal endothelial cells or in response to inflammation. The identification of FCECs during midgestation may be linked to the extensive proliferation of endothelial progenitor cells during corneal morphogenesis.

In summary, our scRNA-Seq analysis of human developing corneas identifies stage-specific definitions of corneal epithelium, stromal and endothelial layers as well as the accessory cell types involved in maintenance of the limbal niche.

## Discussion

Advances in single cell sequencing technologies enable detailed studies of tissues and organs during human development and in adulthood (Behjati et al., 2018). We report here the first comprehensive scRNA-Seq analysis of human cornea encompassing 10-21 PCW and adult samples, providing a detailed map of corneal layer development and cell differentiation. We have used this analysis to identify new markers for qLSCs and TA cells *in vivo*, characterise the changes they undergo during cellular expansion and uncover the transcriptional networks and upstream regulators that maintain qLSCs and TA cell potency. Deposition of our data in open access databases provides a unique opportunity for other researchers to expand this analysis and get new insights on all corneal and conjunctival cell populations.

During development, the eye is constructed from three sources of embryonic precursors: neuroectoderm, surface ectoderm and periocular mesenchyme (Gage et al., 2005). The periocular mesenchyme receives cells from both the neural crest and mesoderm and contributes to multiple mature cell lineages including the corneal endothelium and stroma. Accordingly, our scRNA-Seq showed a large presence of neural crest and mesoderm cells at 10 PCW: these decreased significantly at 12 PCW, coinciding with the expansion of the stromal compartment (keratocytes) and the emergence of blood vessels, melanocytes and corneal nerves. The corneal epithelium itself is formed from bilateral interactions between the neural ectoderm-derived optic vesicles and the cranial ectoderm (Lwigale, 2015). Morphological and ultrastructural studies have shown a segregation between epithelial and stromal cells as early as 6.5 PCW; however, a detailed understanding of conjunctival, corneal and limbal epithelium specification is lacking. Our studies indicate that the first epithelial region to be specified from the ocular surface epithelium is the conjunctiva. This segregation is evident as early as 10 PCW; however, it takes another six weeks until corneal epithelium and the first cluster of proliferating epithelial progenitors appear at 16 PCW. The latter expresses high levels of corneal epithelial progenitor markers (e.g. *KRT15, 14, 19, TP63* among others); nonetheless, none of the qLSCs markers are highly expressed at this stage. This could be due to the lack of limbal niche, which is essential for maintaining the qLSCs population. Although some elements of the niche (e.g. corneal nerves, melanocytes, blood vessels) are present from 12 PCW, the limbal stroma and fibroblasts, that are essential for the generation of necessary ECM and paracrine factor secretion, only become visible at midgestation (20-21 PCW), enabling high expression of some qLSCs markers in the proliferating epithelial progenitor pool (e.g. *TXNIP*), but not all.

In contrast to the rather late segregation of ocular surface epithelium into various subtypes, the corneal endothelium and blood and lymphatic vessels are present as early 12 PCW, with a source of endothelial progenitors established at 16 PCW. Although the endothelial progenitors were present until midgestation, we were unable to locate a similar cluster in adult cornea, which coincides with the inability of corneal endothelium to regenerate (Braunger et al., 2014). It is of interest to note the presence of a different population in adult cornea, identified as FCECs, which are thought to participate in endothelial wound healing under pathological conditions. The FCECs are typical not only of the adult stage, as our single cell analysis also uncovered their presence at midgestation (20-21 PCW) alongside the endothelial progenitor cell cluster. Together these data suggest the establishment of a “reserve endothelial” population at midgestation that may contribute to endothelial wound healing once the endothelial progenitors are no longer present (as in the adult cornea).

The corneal stroma keratocytes expand from 10-18 PCW before consolidating into clusters at midgestation stages. Their development is associated with high expression of key stroma markers including lumican, keratocan and various collagen chains, necessary for the structural integrity of the corneal stroma. The presence of multiple keratocyte clusters can be explained by various states of maturation and proliferation in this compartment as well as their location either at the center or periphery of the cornea. Future studies employing high-resolution spatial transcriptomics based techniques, will enable detailed localisation of these clusters and their developmental maturation. Literature precedence suggests the presence of CSSCs, which are believed to be of neural crest lineage and able to divide extensively *in vitro* and to generate adult keratocytes (Pinnamaneni and Funderburgh, 2012). We were able to identify proliferating progenitors throughout corneal development, however the high expression of cell cycle markers did mask any lineage markers, for this reason we are unsure whether they comprise the CSSCs. Nonetheless, we can detect CSSCs in the adult cornea as a separate cluster associated with high expression of matrix metalloproteinase MMP3 and CD105. There have been extensive discussions in the literature whether CSSCs are bone marrow or neural crest derived. We were unable to observe expression of neural crest markers (*MITF, PAX6, SOX9* etc) in the CSSCs cluster; however, our analysis did uncover a population of neural crest progenitors (LNCPs) located under the limbal crypts next to limbal basal epithelium, expressing neural crest markers (*PAX6, MITF*) and appreciable levels of *Ki67*. Together these findings suggest that CSSCs and LNCPs are two different cell populations, with the first most likely to contribute to the stroma regeneration and the second to the maintenance of qLSCs.

The renewal of corneal epithelium is critically reliant on LSCs. Loss of LSCs results in limbal stem cell deficiency (LSCD) characterised by corneal neovascularization, chronic inflammation stromal scarring, corneal opacity and loss of vision (Baylis et al., 2013). Transplantation of autologous *ex vivo* expanded LSCs from a healthy contralateral eye onto the patient’s damaged eye is an established treatment for patients with total/severe unilateral LSCD (Holoclar, European Medicines Agency, Kolli et al. 2009). The frequency of stem cells in the transplanted cell population is a key success factor for restoration of vision (Rama et al., 2010); hence identification, quantitation and enrichment of LSCs prior to transplantation is an important and yet unmet area for research. Several putative LSCs markers have been identified including ΔNp63α, ABCG2, ABCB5, C/EBPσ, Bmi1, Notch-1 amongst others. (Mort et al., 2012). To verify these findings, we took an unbiased approach by identifying highly expressed marker genes found in qLSCs cluster relative to other clusters. We found five new marker genes not previously associated with qLSCs, namely *GPHA2, CASP14, MMP10, MMP1* and *AC093496.1 (*Lnc-XPC-2). Three of these, GPHA2, MMP10 and MMP1 were localised to the limbal crypts, overlapping in expression with each other or KRT15 and ΔNp63 but not the proliferation marker, Ki67. Downregulation of *GPHA2* using RNAi greatly reduced colony forming efficiency, obliterated holoclones and induced differentiation. GPHA2 has recently been identified as a qLSCs marker in the mouse cornea (Ruby Shalom-Feuerstein, personal communication), further corroborating data presented in this manuscript. Until the advent of single cell sequencing, marker identification relied on differential gene expression between LECs and differentiated cells; however, the *ex vivo* expanded LEC cultures are enriched for progenitors; hence, the identified markers most likely represent progenitor cells markers, which explain why the qLSCs markers identified in this study have not been reported previously.

Given the importance of *ex vivo* LECs expansion for LSCD treatment, we used single cell sequencing to compare cells before and after *ex vivo* expansion under two different culture conditions: 3T3 feeders and HAM. Although no significant differences were observed between the two expansion methods, the *ex vivo* LECs showed a significant downregulation of the qLSCs markers identified in this study (e.g. *GPHA2*) and acquisition of markers associated with proliferative limbal epithelial progenitors. The same phenomenon was observed in limbal dysplasia *in vivo*, suggesting that this change in transcriptional profile could be due to several events including stimulation of proliferation, which brings the LSCs out of their quiescent state, as well as the incomplete replication of limbal niche under *ex vivo* culture conditions.

Our integrated single cell analysis revealed very close and well-balanced interactions between qLSCs, immune cells, blood cells and corneal nerves. These findings are replicated from single cell studies in the mouse limbus (Ruby Shalom-Feuerstein, personal communication) and corneal immune related pathologies (Steven-Johnson syndrome, chronic limbitis or ocular pemphigoid) where excess ocular surface inflammation involving the limbus can lead to LSCD. Together these data suggest the need for a complete replication of the limbal niche during *ex vivo* propagation methods in order to fully support qLSCs survival, self-renewal and potency before clinical cell based interventions. An interesting observation is that *ex vivo* expanded LECs with the current methods (HAM or 3T3 feeders) confer an overall 75% clinical success rate in cultured LEC transplantation. This suggests that *ex vivo* expanded LECs may regain their gene expression profile akin to qLSCs upon re-establishing interactions with the remaining limbal niche *in vivo*.

Finally, we were also interested to validate the application of single cell sequencing as a platform for gaining insights into corneal diseases. We focused on keratoconus, a corneal disorder characterized by progressive thinning and changes in the shape of the cornea, which affects approximately 1 in 2,000 individuals worldwide. To date there have been several proteomic studies of corneal stroma from affected patients (Chaerkady et al., 2013; Shinde et al., 2019; Vallabh et al., 2017; Wojcik et al., 2013), revealing EIF2 and mTor signalling, oxidative phosphorylation, and mitochondrial dysfunction to be at the heart of molecular changes in these patients. These same pathways were identified from the scRNA-Seq of only two patients and one unaffected subject. Importantly, we were able to identify the reason behind corneal epithelial thinning which is due to differentiation of TA cells towards superficial cells, resulting in a decreased pool of progenitors that are able to replenish the central corneal epithelium. These findings when validated in a larger number of patients, may have important implications for future treatment of these patients.

In summary, the single cell analysis described herein provides the first cell type specific information for all the cells and layers found in the adult human cornea, limbus and surrounding conjunctiva. By expanding this analysis to the developmental cornea and conjunctiva samples obtained from 10-21 PCW, we were able to determine the stage specific definitions of conjunctival, corneal and progenitor epithelial cells populations, establishment of the limbal niche as well as segregation of stroma and endothelium and their associated progenitor cell subpopulations. Bioinformatic comparison of adult cell clusters identified GPHA2, a novel cell-surface marker for quiescent limbal stem cells (qLSCs), whose function is to maintain qLSCs self-renewal. Our results also provide an excellent platform for using scRNA-Seq to compare the *ex vivo* expanded LECs to their native counterparts and comparing two different widely used cell expansion methods. Analysis of these datasets indicated that culture methods do not account for the transcriptional differences observed between the expanded limbal epithelial cells and qLSCs *in vivo*. The very close interactions between qLSCs and the different elements of the niche (immune cells, blood cells, corneal nerves, limbal fibroblasts and stroma) bring to the forefront the importance of limbal niche in maintaining the qLSCs quiescence and potency. Overall, the data presented herein, showcase the ability of scRNA- and ATAC-Seq to assess multiple datasets from the developmental and adult cornea, particularly the limbus under normal and disease states in a comprehensive manner, which will help to define pathways/genes that could lead to improvement in *ex vivo* expansion methods for cell based replacement therapies for corneal disease and repair of the limbal niche before or as part of clinical cell based interventions.

## Acknowledgments

We are grateful to MRC UK for funding this work (MR/S035826/1), the Newcastle University Flow Activated Cell Sorting Facility for help with cell sorting and Dr. Dean Hallam for drawing the graphical abstract. The human embryonic and fetal material was provided by the joint MRC/Wellcome Trust Human Developmental Biology Resource (MR/R006237/1). The adult eye tissue was provided by NHS Blood and Transplant Tissue and Eye Services following ethical approval (18/YH/04/20).

## Author contributions

JC: setup and optimisation of scRNA- and ATAC-Seq and tissue dissociations, performed experiments, experimental design, data collection and data submission

RQ: setup bioinformatics pipeline, performed bioinformatics analysis, figure preparation and data submission

DZ: performed IHC experiments, data collection, IHC data analysis, IHC figure preparation NM, MMM, SB, CY, GR, and JMC: performed experiments, data collection

GR, MH: data analysis

SL, DH: facilitated embryonic and fetal sample collection for the study

AJ, PR, SG: facilitated adult sample collection for the study

CC, FF: experimental design and fund raising

LA: performed experiments, data collection, data analysis, figure preparation, experimental design and manuscript writing

ML: designed and performed experiments, data collection and analysis, figure preparation, manuscript writing and fund raising.

All authors contributed and approved the final version of the manuscript.

## Notes

### Competing Interest Statement

The authors have declared no competing interest.

