## Supplementary material for "A single cell atlas of human cornea that defines its development, limbal stem and progenitor cells and the interactions with the limbal niche": Suppl. Figures and Methods

#### Inventory of supplemental Information

##### Supplemental Figures and Legends

Figure S1, related to Figure 1

Figure S2, related to Figure 1

Figure S3, related to Figure 1

Figure S4, related to Figure 1

Figure S5, related to Figure 1

Figure S6, related to Figure 1

Figure S7, related to Figure 1

Figure S8, related to Figure 1

Figure S9, related to Figure 1

Figure S10, related to Figure 1

Figure S11, related to Figure 1

Figure S12, related to Figure 1

Figure S13, related to Figure 1

Figure S14, related to Figure 1

Figure S15, related to Figure 4

Figure S16, related to Figure 4

Figure S17, related to Figure 5

Figure S18, related to Figure 6

Figure S19, related to Figure 6

Figure S20, related to Figure 7

Figure S21, related to Figure 7

**Table S1**: A full list of highly and differentially expressed genes between the 21 clusters identified in the human cornea and surrounding conjunctiva corresponding to Figure S1.

**Table S2**: A full list of highly and differentially expressed genes between the immune cell clusters identified in the human cornea and surrounding conjunctiva corresponding to Figure S11A.

**Table S3**: A full list of differentially accessible (DA) peaks, enhancers and enriched motif for qLSCs cluster 9 comparison versus all epithelial cells identified in the adult cornea and the surrounding conjunctiva.

**Table S4**: A full list of differentially accessible (DA) peaks, enhancers and enriched motif for TA cluster 4 comparison versus all epithelial cells.

**Table S5:** List of significant regulators of gene expression in qLSCs (cluster 9) and TA cells (cluster 4).

**Table S6**: A list of significant ligand-receptor interactions between qLSCs (cluster 9) and immune cells (clusters 15 and 17) revealed by CellPhoneDB.

**Table S7**: A full list of highly and differentially expressed genes between the three additional clusters identified in the *ex vivo* expanded LECs corresponding to Figure 5A.

**Table S8**: A full list of differentially expressed genes between the additional clusters 1 and 2 found in the *ex vivo* expanded LECs and qLSCs (cluster 9) and the basal conjunctival epithelial (cluster 0) of adult cornea and conjunctiva respectively.

**Table S9**: A full list of highly and differentially expressed genes between the original 21 clusters found in the adult cornea and conjunctiva and the six additional clusters identified in the cornea with limbal dysplasia corresponding to Figure 6B.

**Table S10**: A full list of highly and differentially expressed genes between the original 21 clusters found in the adult cornea and conjunctiva and the Keratoconus cornea samples corresponding to Figure 7A.

**Table S11**: A full list of differentially expressed genes in stroma keratocytes (cluster 12) and TA cells (cluster 4) between Keratoconus cornea patients and unaffected subjects.

**Table S12:** A list of signalling pathways enriched in the Keratoconus stroma keratocytes (cluster 12) and TA (cluster 4) generated by IPA.

**Table S13**: A list of significant regulators of gene expression in the Keratoconus stroma keratocytes (cluster 12) generated by IPA.

**Table S14:** A full list of highly and differentially expressed genes between clusters identified in scRNA-Seq of embryonic and fetal cornea and conjunctiva samples from 10-21 PCW corresponding to Figure 8.

**Table S15**: List of antibodies used for the IHC.

**Table S16:** List of primers used for the qRT-PCR analysis.

#### Supplemental Experimental Procedures

#### Supplemental Reference

#### Supplemental Figures and Legends

### Figure S1

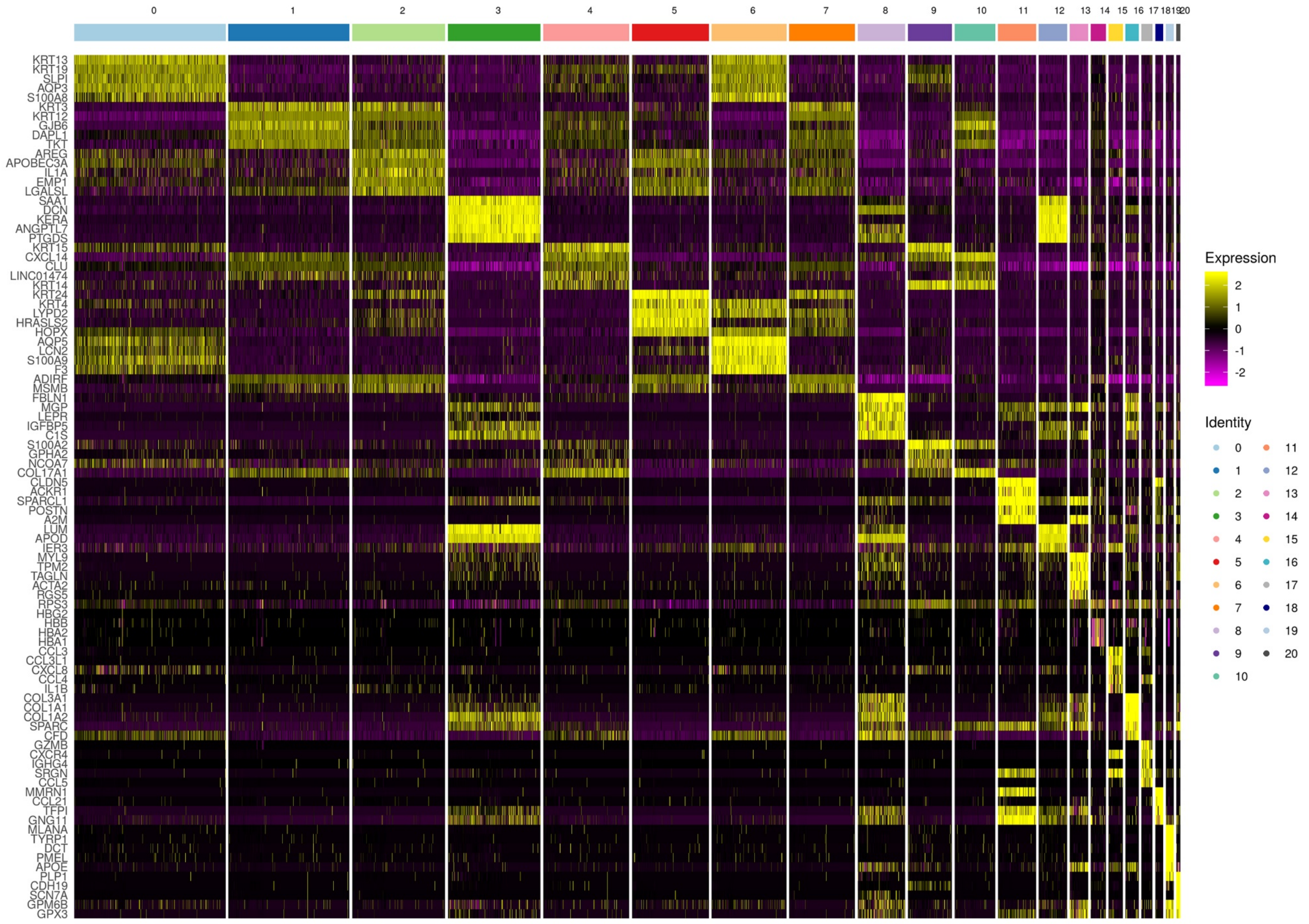
0 1 2 3 4 5 6 7 8 9 10 11 12 13 14 15 16 17 18 19 20
KRT13
KRT19
SLPI
AQP3
S100A8
KRT3
KRT12
GJB6
DAPL1
TKT
AREG
APOBEC3A
IL1A
EMP1
LGALSL
SAA1
DCN
KERA
ANGPTL7
PTGDS
KRT15
CXCL14
CLU
LINC01474
KRT14
KRT24
KRT4
LYPD2
HRASLS2
HOPX
AQP5
LCN2
S100A9
F3
ADIRF
MSMB
FBLN1
MGP
LEPR
IGFBP5
C1S
S100A2
GPHA2
NCOA7
COL17A1
CLDN5
ACKR1
SPARCL1
POSTN
A2M
LUM
APOD
IER3
MYL9
TPM2
TAGLN
ACTA2
RGS5
RPS3
HBG2
HBB
HBA2
HBA1
CCL3
CCL3L1
CXCL8
CCL4
IL1B
COL3A1
COL1A1
COL1A2
SPARC
CFD
GZMB
CXCR4
IGHG4
SRGN
CCL5
MMRN1
CCL21
TFPI
GNG11
MLANA
TYRP1
DCT
PMEL
APOE
PLP1
CDH19
SCN7A
GPM6B
GPX3
Expression
2
1
0
-1
-2
Identity

**Figure S1: Clustering analysis reveals the presence of 21 clusters in the adult human cornea and surrounding conjunctiva** (see also Figure 1 and Table S1). The top 5 markers used for cluster annotation are shown with a complete list of differentially expressed genes in Table S1.

**Figure S2**

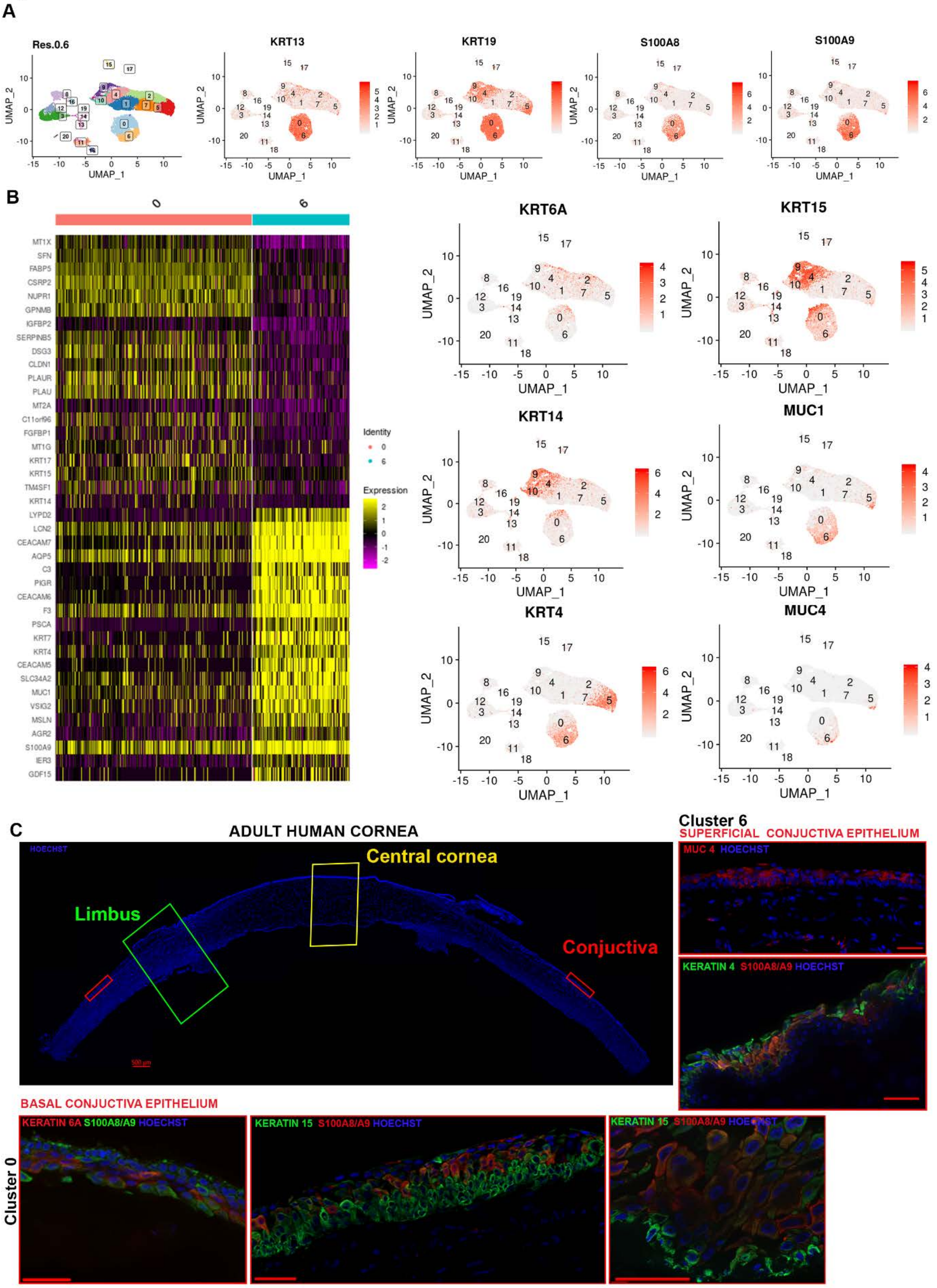
A
Res.0.6
KRT13
KRT19
S100A8
S100A9
UMAP_1
UMAP_2
B
0
6
MT1X
SFN
FABP5
CSRP2
NUPR1
GPNMB
IGFBP2
SERPINB5
DSG3
CLDN1
PLAUR
PLAU
MT2A
C11orf96
FGFBP1
MT1G
KRT17
KRT15
TM4SF1
KRT14
LYPD2
LCN2
CEACAM7
AQP5
C3
PIGR
CEACAM6
F3
PSCA
KRT7
KRT4
CEACAM5
SLC34A2
MUC1
VSIG2
MSLN
AGR2
S100A9
IER3
GDF15
Identity
Expression
KRT6A
KRT15
KRT14
MUC1
KRT4
MUC4
C
ADULT HUMAN CORNEA
HOECHST
Central cornea
Limbus
Conjuctiva
Cluster 6
SUPERFICIAL CONJUCTIVA EPITHELIUM
MUC 4 HOECHST
KERATIN 4 S100A8/A9 HOECHST
BASAL CONJUCTIVA EPITHELIUM
Cluster 0
KERATIN 6A S100A8/A9 HOECHST
KERATIN 15 S100A8/A9 HOECHST
KERATIN 15 S100A8/A9 HOECHST

**Figure S2: Characterisation of the basal and superficial conjunctival epithelial clusters** (see also Figure 1 and Table S1). **A**) UMAP of adult human cornea and conjunctiva with superimposed single gene expression plots showing the expression of four highly expressed markers in both basal and superficial conjunctival epithelium; **B**) Comparative heatmap showing differentially expressed genes between clusters 0 and 6 on the left with representative gene expression plots of markers with higher expression in cluster 0 (*KRT6A, KRT15, KRT14*) or cluster 6 (*MUC1, KRT4, MUC4*) on the right hand side panel; **C**) Immunohistochemical analysis showing the expression of KRT15 and KRT6A predominantly in the basal conjunctival epithelium and KRT4 and MUC4 in the superficial conjunctival epithelium. S100A8/A9 is expressed throughout the conjunctival epithelium, Scale bars: 50 µm.

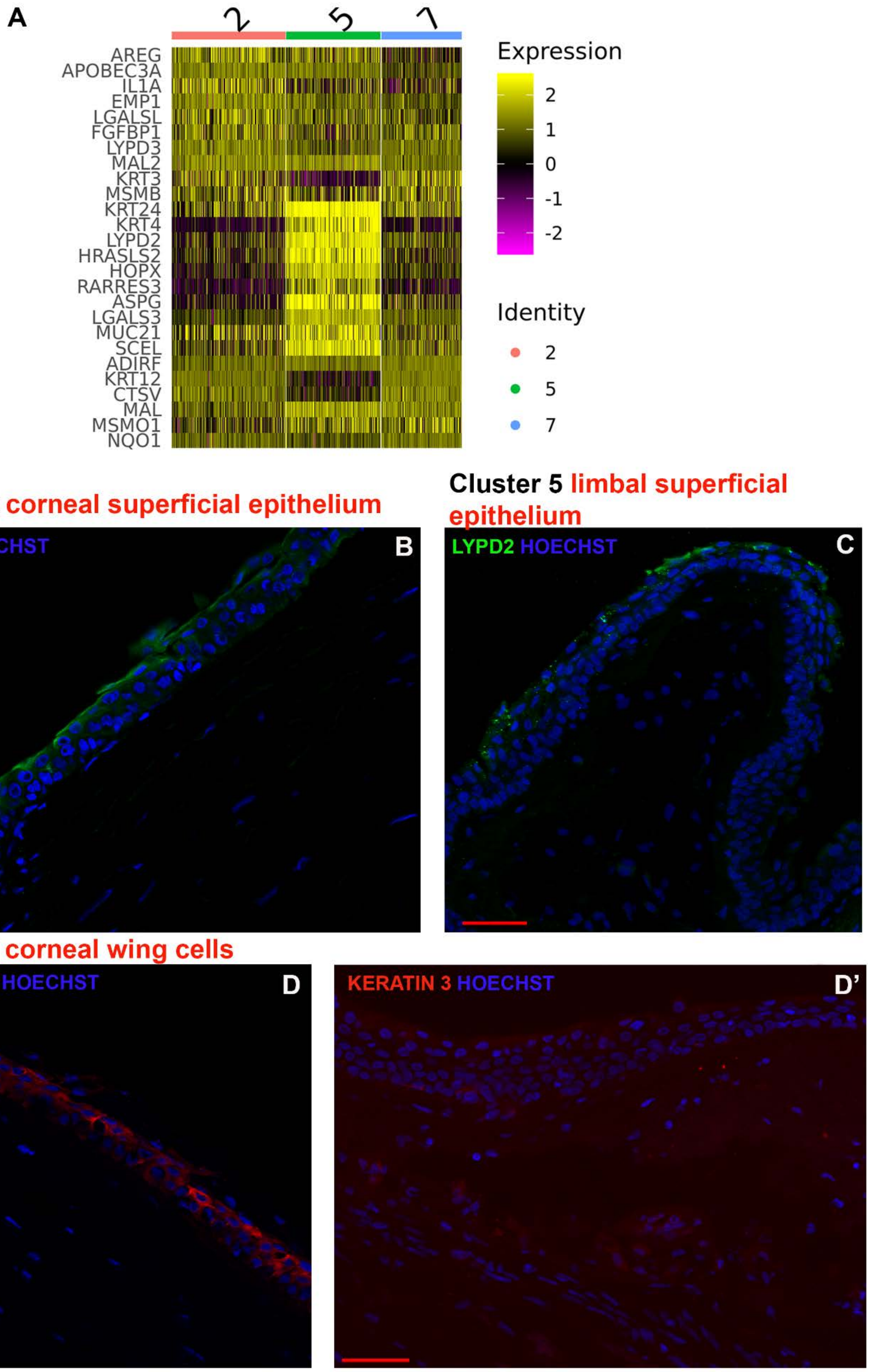
A
2
5
7
AREG
APOBEC3A
IL1A
EMP1
LGALSL
FGFBP1
LYPD3
MAL2
KRT3
MSMB
KRT24
KRT4
LYPD2
HRASLS2
HOPX
RARRES3
ASPG
LGALS3
MUC21
SCEL
ADIRF
KRT12
CTSV
MAL
MSMO1
NQO1
Expression
2
1
0
-1
-2
Identity
2
5
7
Cluster 2 corneal superficial epithelium
Cluster 5 limbal superficial epithelium
AREG HOECHST
B
LYPD2 HOECHST
C
Cluster 7 corneal wing cells
KERATIN 3 HOECHST
D
KERATIN 3 HOECHST
D’

**Figure S3**

**Figure S3: Characterisation of the corneal and limbal superficial epithelium** (see also Figure 1 and Table S1). **A**) Comparative heatmap showing differentially expressed genes between clusters 2, 5 and 7; **B-D**) IHC analysis showing AREG (**B**), LYPD2 (**C**) and KRT3 (**D**) expression in the corneal superficial epithelium, limbal superficial epithelium and corneal wing cells respectively. **D'** shows the lack of KRT3 expression in the limbal epithelium. Scale bars: 50 µm.

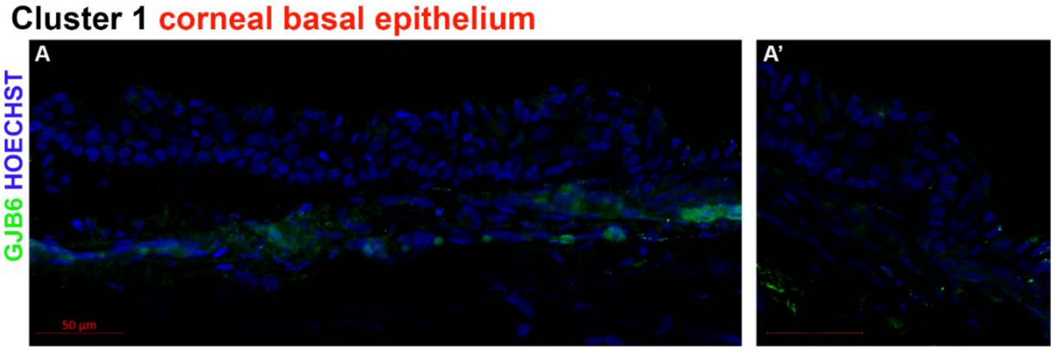

**Figure S4**

**Figure S4: Characterisation of the corneal basal epithelium** (see also Figure 1 and Table S1). **A**) IHC analysis showing the expression of GJB6 in the basal layer of the corneal epithelium; A' shows very low GJB6 expression in the limbal basal epithelium. Scale bars: 50µm.

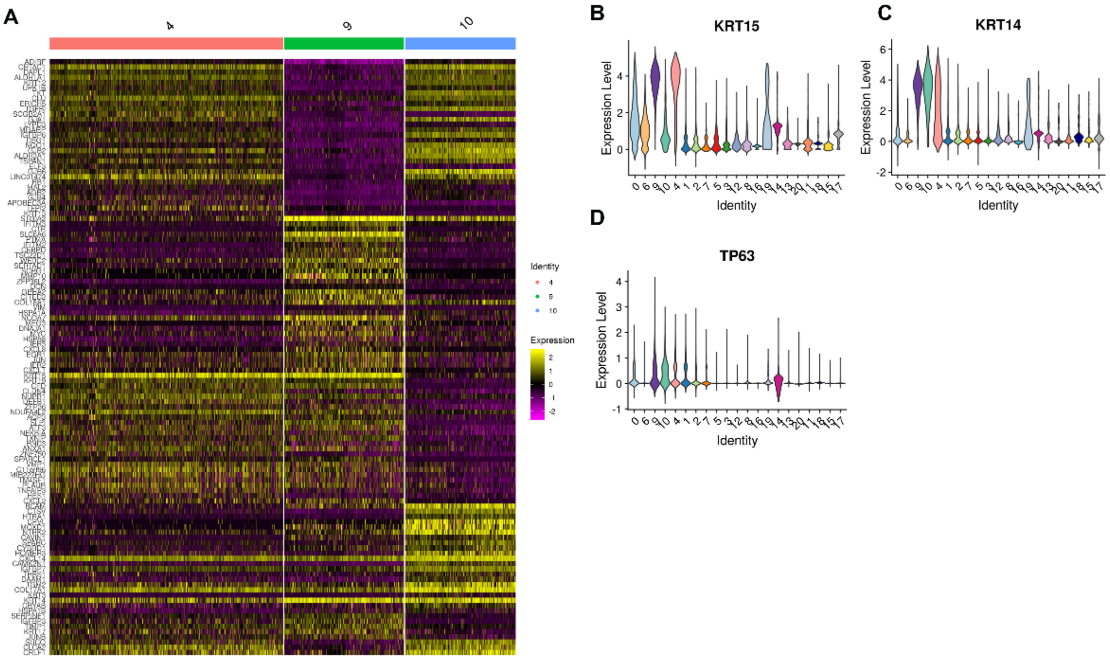

**Figure S5**

**Figure S5**: **Detailed transcriptional analysis of clusters 4, 9 and 10** (see also Figure 1 and Table S1). **A**) Comparative gene expression heatmap between clusters 4, 9 and 10; **B-D**) Violin plots showing expression of *KRT15* (**B**), *KRT14* (**C**) and *TP63* (**D**).

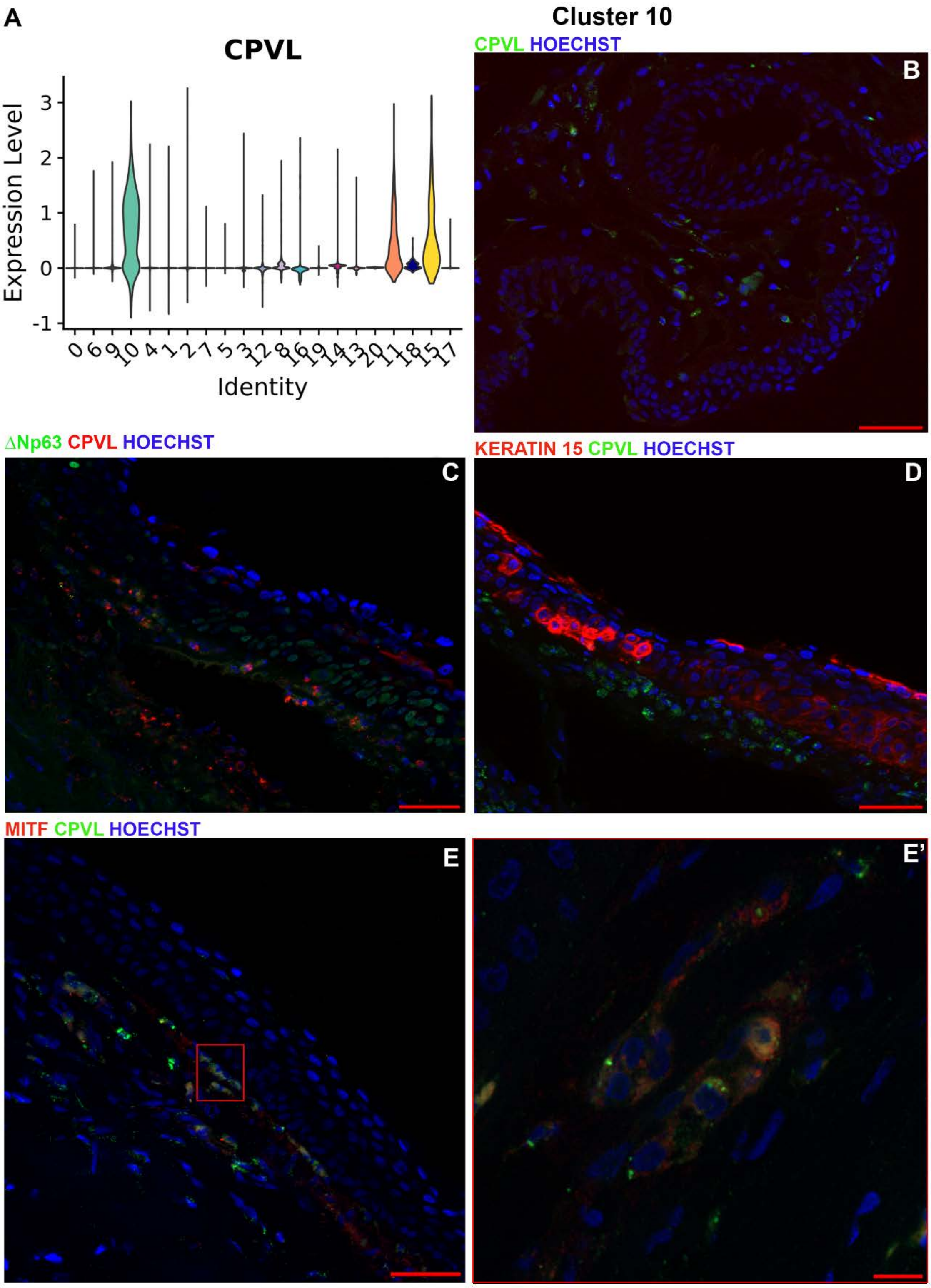

**Figure S6**

**Figure S6: Detailed analysis of cluster 10 representing the limbal neural crest progenitor cells** (see also Figure S5 and Table S1). **A**) Violin plot showing the highest *CPLV* expression in cluster 10; **B**) IHC analysis showing CPVL expression adjacent to the basal limbal epithelium; **C**) Combined CPVL and ΔNp63 IHC analysis showing a small minority of co-expressing cells; **D**) IHC analysis showing the location of CPVL immunopositive cells next to the basal limbal epithelial cells marked by KRT15 expression; **E**, **E'**) Co-expression of CPVL with MITF in stromal cells located next to the limbal basal epithelium. Red boxed inset shows the higher magnification of CPVL and MITF co-expression cells (**E'**). Scale bars: 50 µm (B, C, D, E) and 10 µm (E').

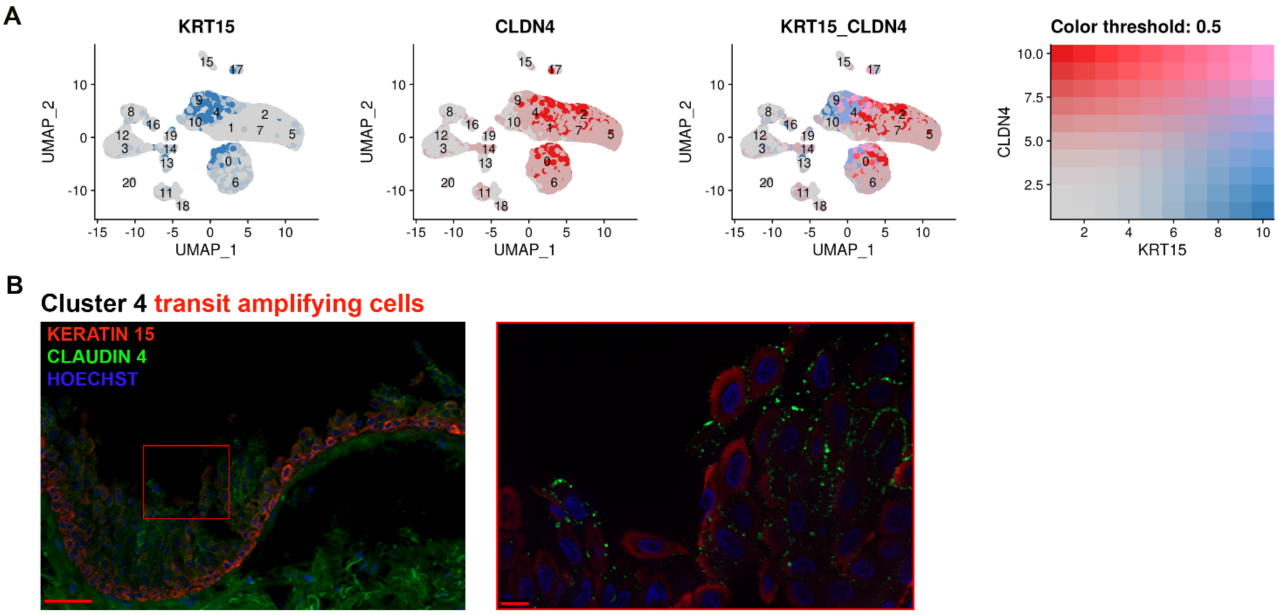

**Figure S7**

**Figure S7: Detailed characterisation of cluster 4** (see also Figure S5 and Table S1). **A**) *KRT15* and *CLDN4* UMAP superimposed expression plots showing co-localisation of these two genes in cluster 4; **B**) IHC analysis showing the co-expression of KRT15 and CLDN4 in the cells that are migrating away from the basal limbal epithelium. Red inset: high magnification image showing the double positive cells for $KRT15^+$ and $CLDN4^+$ in the area indicated in B. Scale bars: 50 µm and 10 µm (red inset).

**Figure S8**

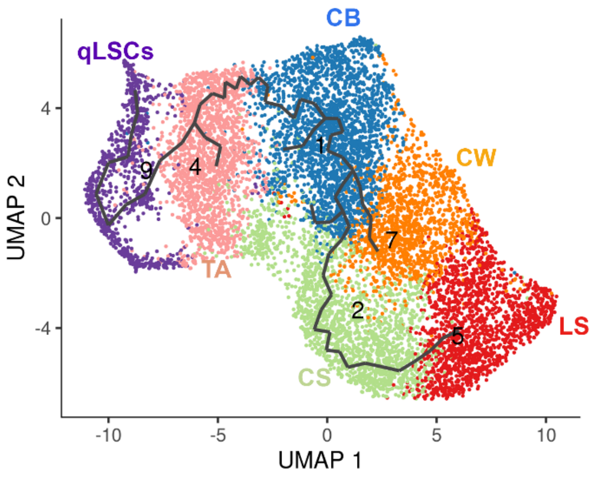

**Figure S8: Pseudotime analysis of stem, progenitor and differentiated corneal epithelial cell clusters** (see also Figure 1 and Table S1).

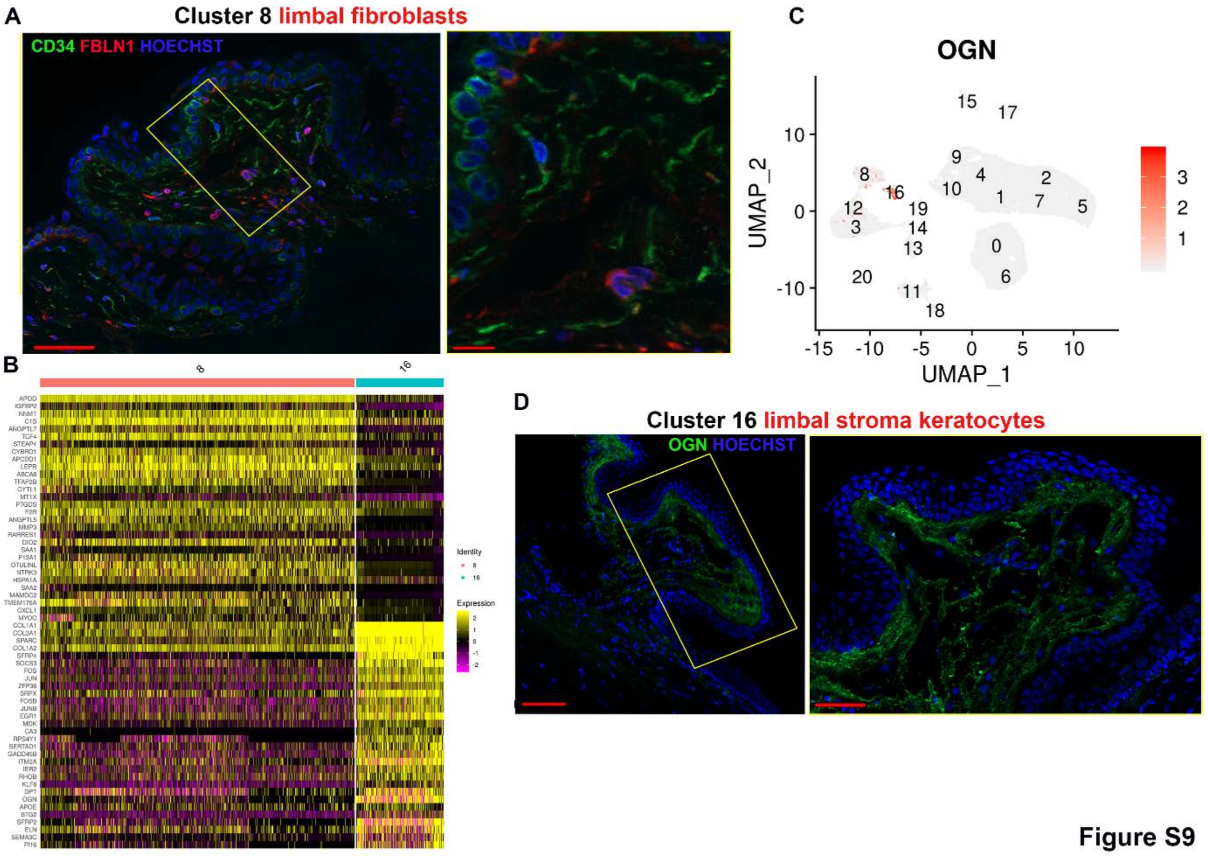

**Figure S9**

**Figure S9: Detailed characterisation of cluster 8 and 16** (see also Figure 1 and Table S1). **A**) IHC analysis showing expression of FBLN1 underneath the limbal crypts. No co-expression with CD34 was observed. The yellow inset shows a high magnification of IHC on the right hand side of the image; **B**) Comparative expression heatmap showing the differentially expressed genes between clusters 8 and 16; **C**) *OGN* expression superimposed over the UMAP showing high and predominant expression in cluster 16; **D**) IHC analysis showing expression of OGN in the limbal stroma. Since the stroma ECM is secreted by the keratocytes, this cluster was defined as limbal stroma keratocytes. The yellow inset shows a high magnification of IHC on the right hand side of the image. Scale bars: 50 µm and 10 µm (yellow insets).

### Figure S10

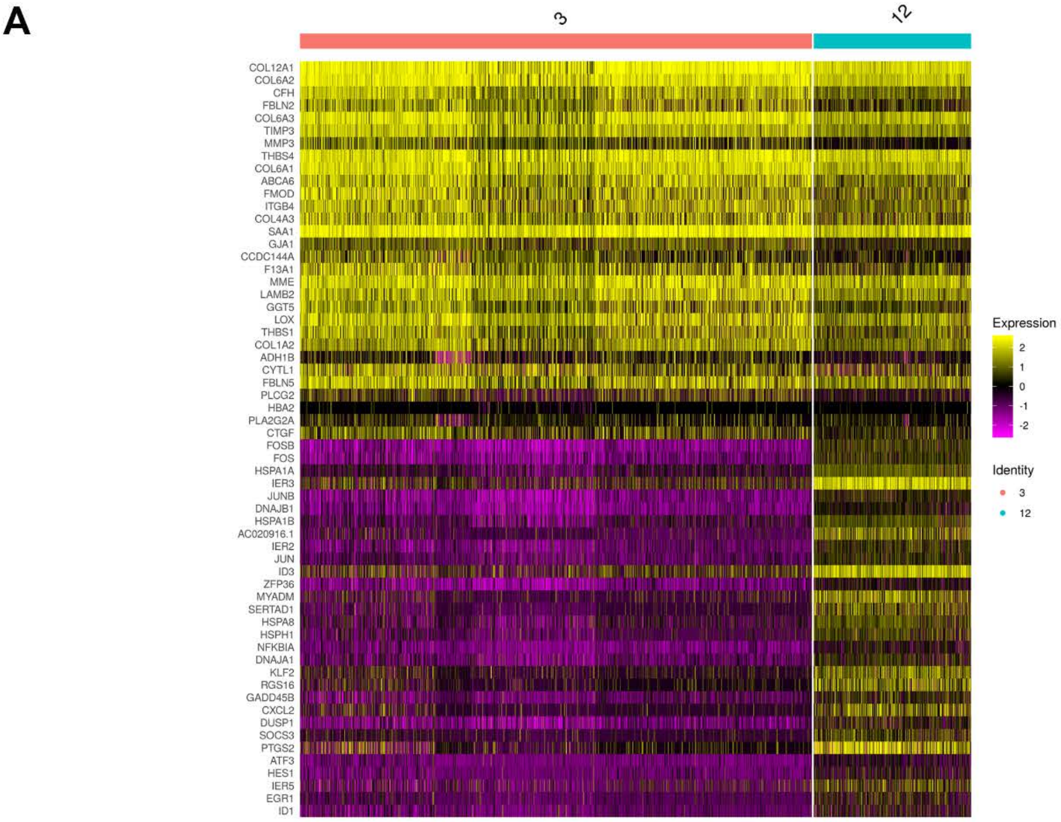

**Cluster 3 corneal stromal stem cells**

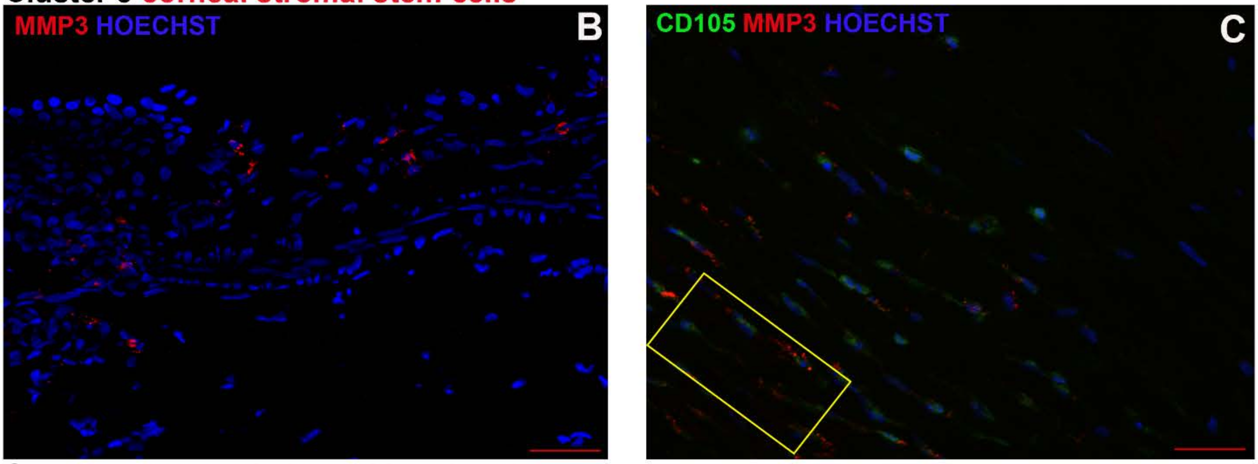

**Cluster 12 corneal stromal keratocytes**

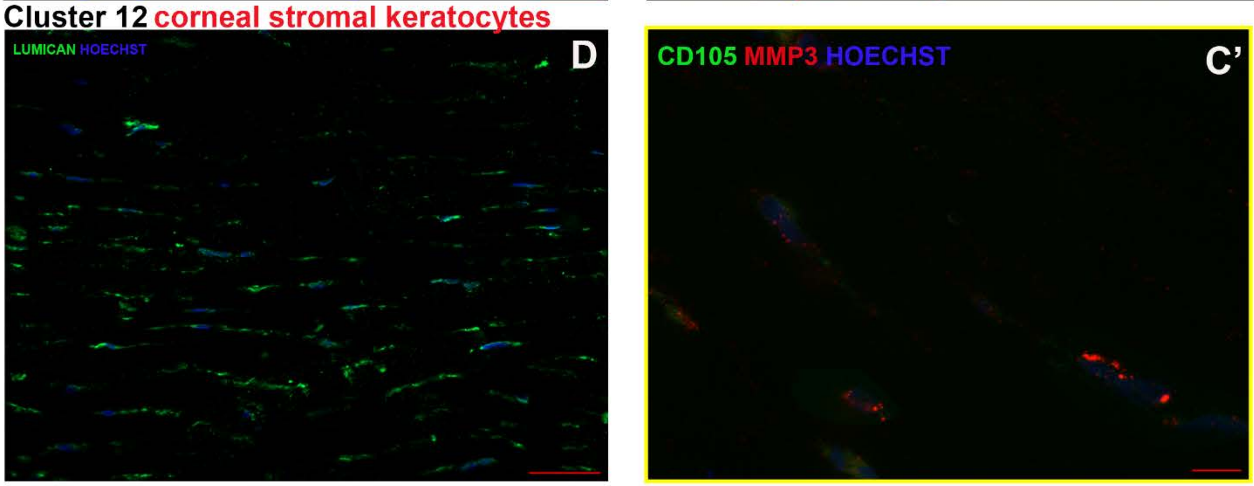

**Figure S10: Detailed characterisation of clusters 3 and 12** (see also Figure 1 and Table S1). **A**) Comparative heatmap showing the differentially expressed genes between clusters 3 and 12; **B-C**) IHC analysis showing the expression of MMP3 in the peripheral and central cornea and co-expression with CD105. The yellow inset shows a high magnification of IHC in **C'**; **D**) Lumican expression in the stroma of central cornea. Since the stroma ECM is secreted by the keratocytes, cluster 12 was defined as central stroma keratocytes. Scale bars: 50 µm (A, B, C, D) and 10 µm (C').

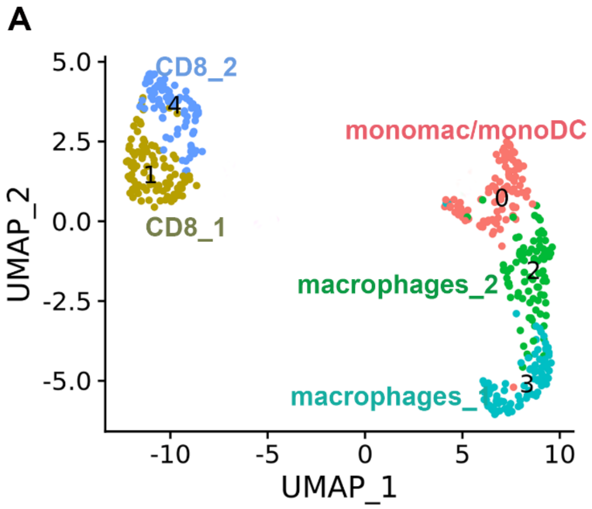

**Figure S11**

**Figure S11: Further characterisation of immune cells (cluster 15 and 17) found in the adult human cornea and conjunctiva** (see also Figure 1 and Table S2). **A**) UMAP showing the presence of CD8, macrophages and dendritic cells. A manual doublet discrimination step was performed to remove a small cell cluster which expressed both epithelial and immune cells markers.

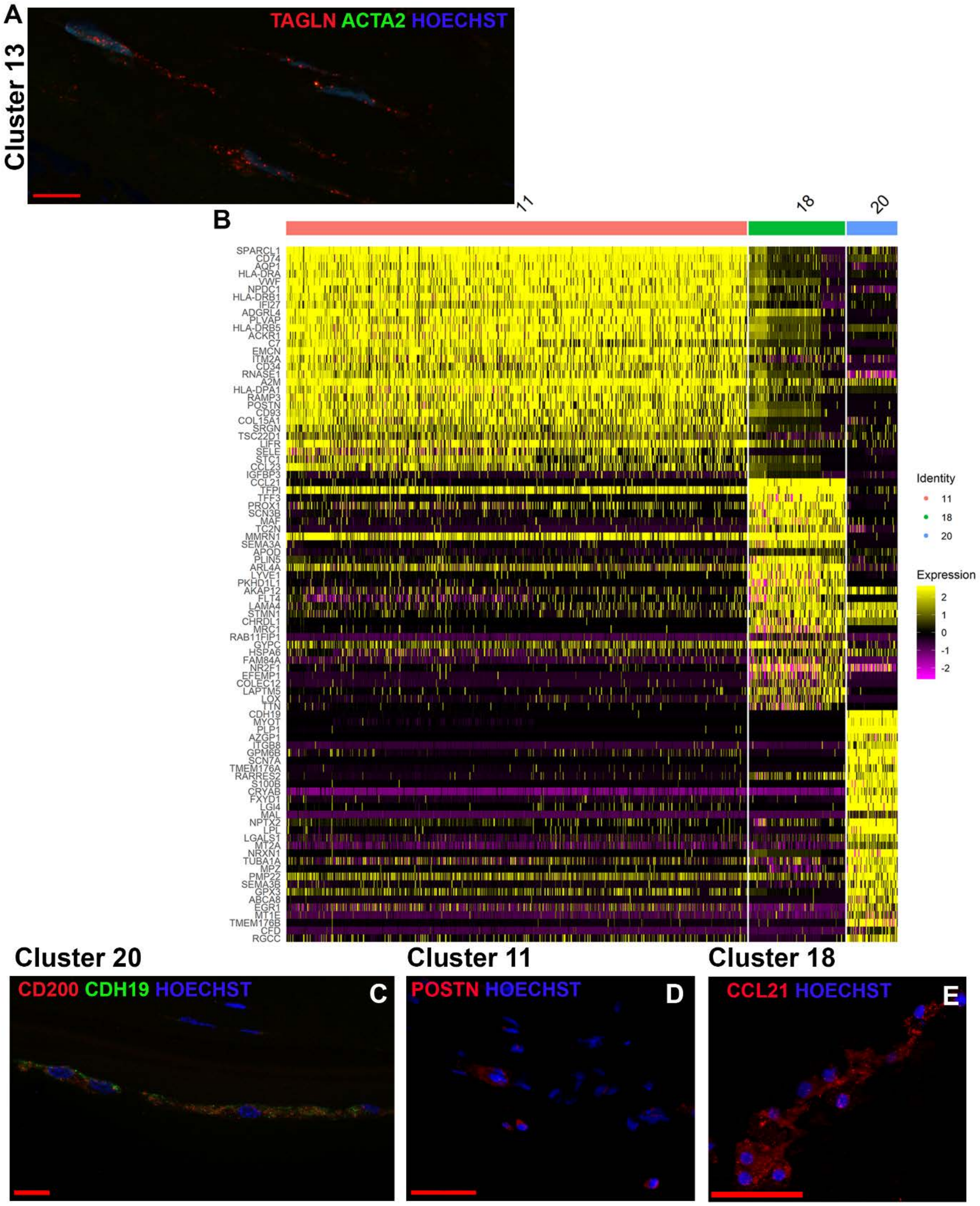

**Figure S12**

**Figure S12: Detailed characterisation of endothelial cell clusters** (see also Figure 1 and Table S1). **A**) Co-expression of TAGLN and ACTA2 in fibroblastic corneal endothelial cells, located in the stroma of limbal and peripheral cornea; **B**) Comparative heatmap showing the differentially expressed genes between clusters 11, 18 and 20; **C**) IHC analysis showing the expression of CDH19 (and co-expression with CD200) in the corneal endothelium; **D**) IHC analysis showing the expression of POSTN in the limbal blood vessels; **E**) IHC analysis showing the expression of CCL21 in lymphatic vessels found in the limbus and conjunctiva. Scale bars: 50 µm (D, E,) and 10 µm (A, C).

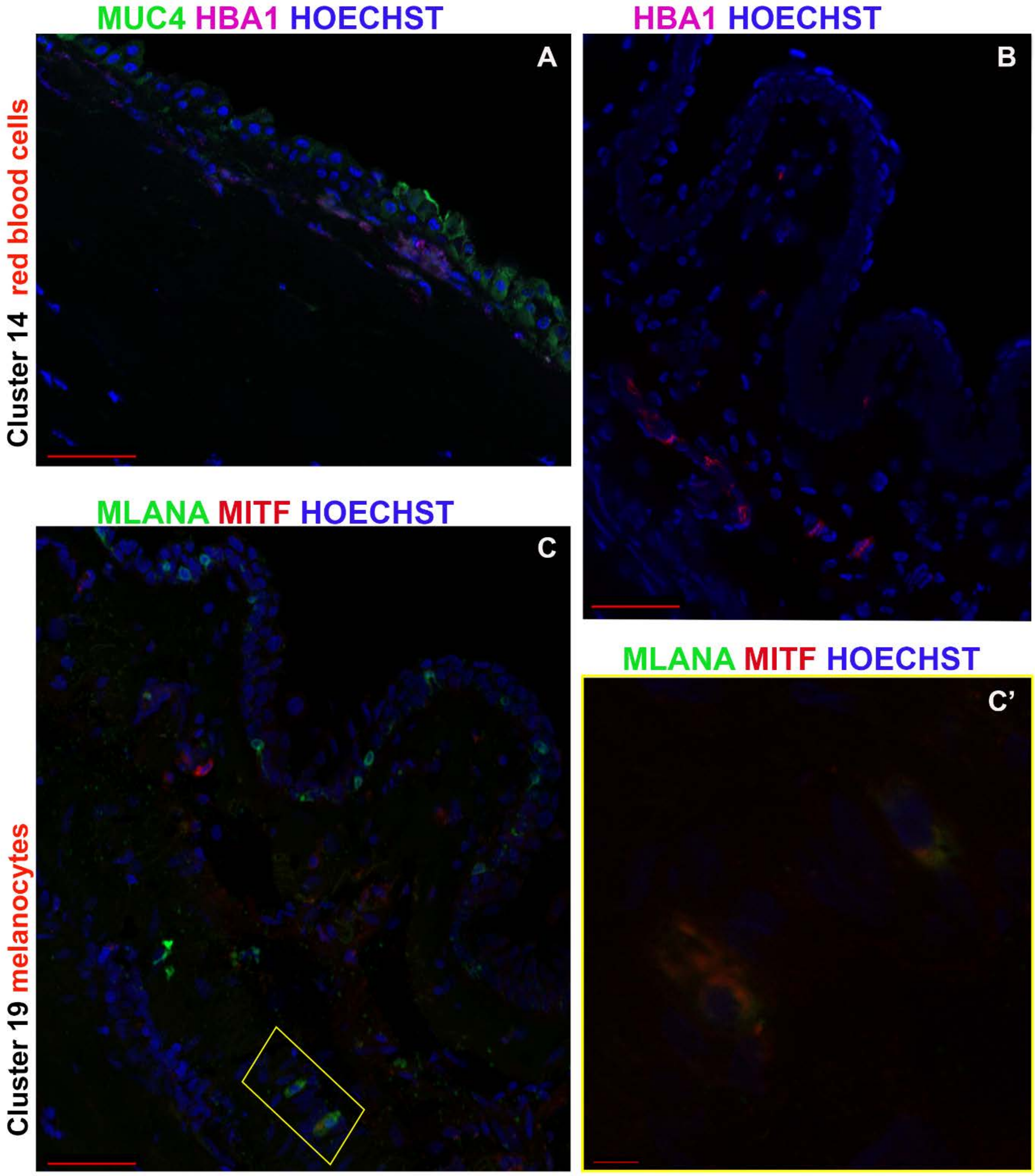

Figure S13

**Figure S13: Further characterisation of red blood cells and melanocytes** (see also Figure 1 and Table S1). **A, B**) IHC analysis showing the presence of HBA1+ red blood cells under the conjuctival epithelium marked by MUC4 staining (**A**) as well scattered under the limbal crypts (**B**); **C**) Co-expressing MLANA and MITF melanocytes are found underneath the limbal crypt. **C'**) The yellow inset shows a high magnification of IHC. Scale bars: 50 µm (A, B, C) and 10 µm (C').

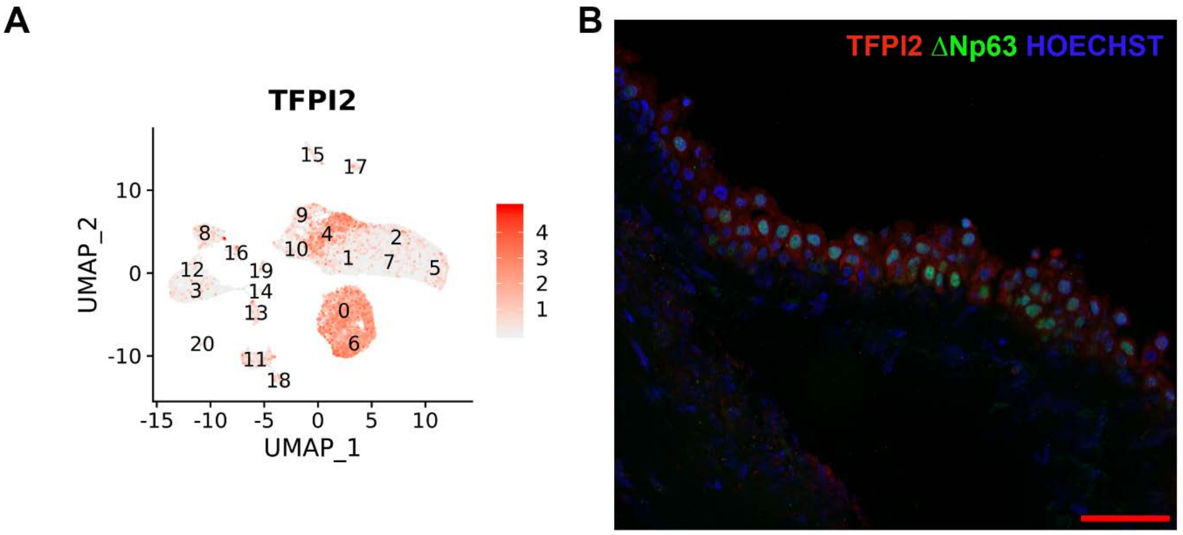

**Figure S14**

**Figure S14: TFPI2 expression in TA cells** (see also Figure 1 and Table S1). **A**) High TFPI2 expression in TA cells (cluster 4) shown through a gene expression map superimposed to the UMAP; **B**) Co-localisation of TFPI2 and ΔNp63 in TA cells. Scale bars: 50 µm.

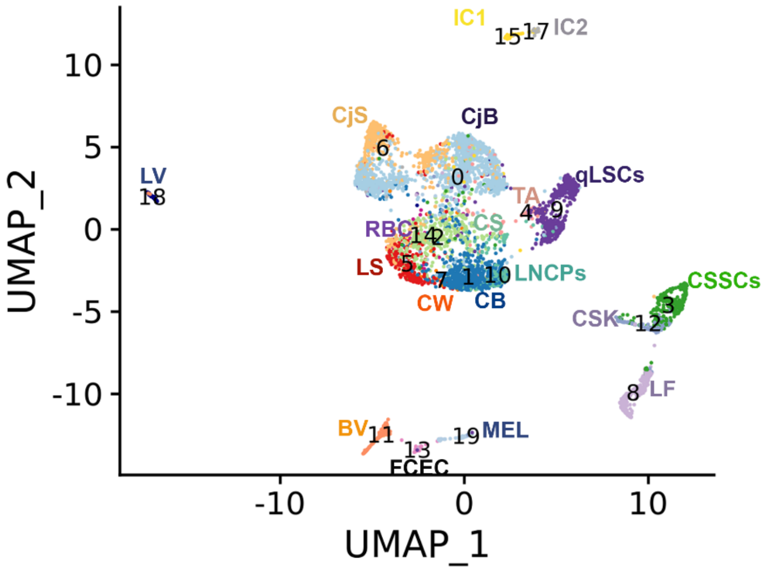

**Figure S15**

**Figure S15: UMAP of scATAC-Seq of human adult cornea and conjunctiva** (see also Figure 4). All cluster annotations are the same as in Figure 1A.

**Figure S16**

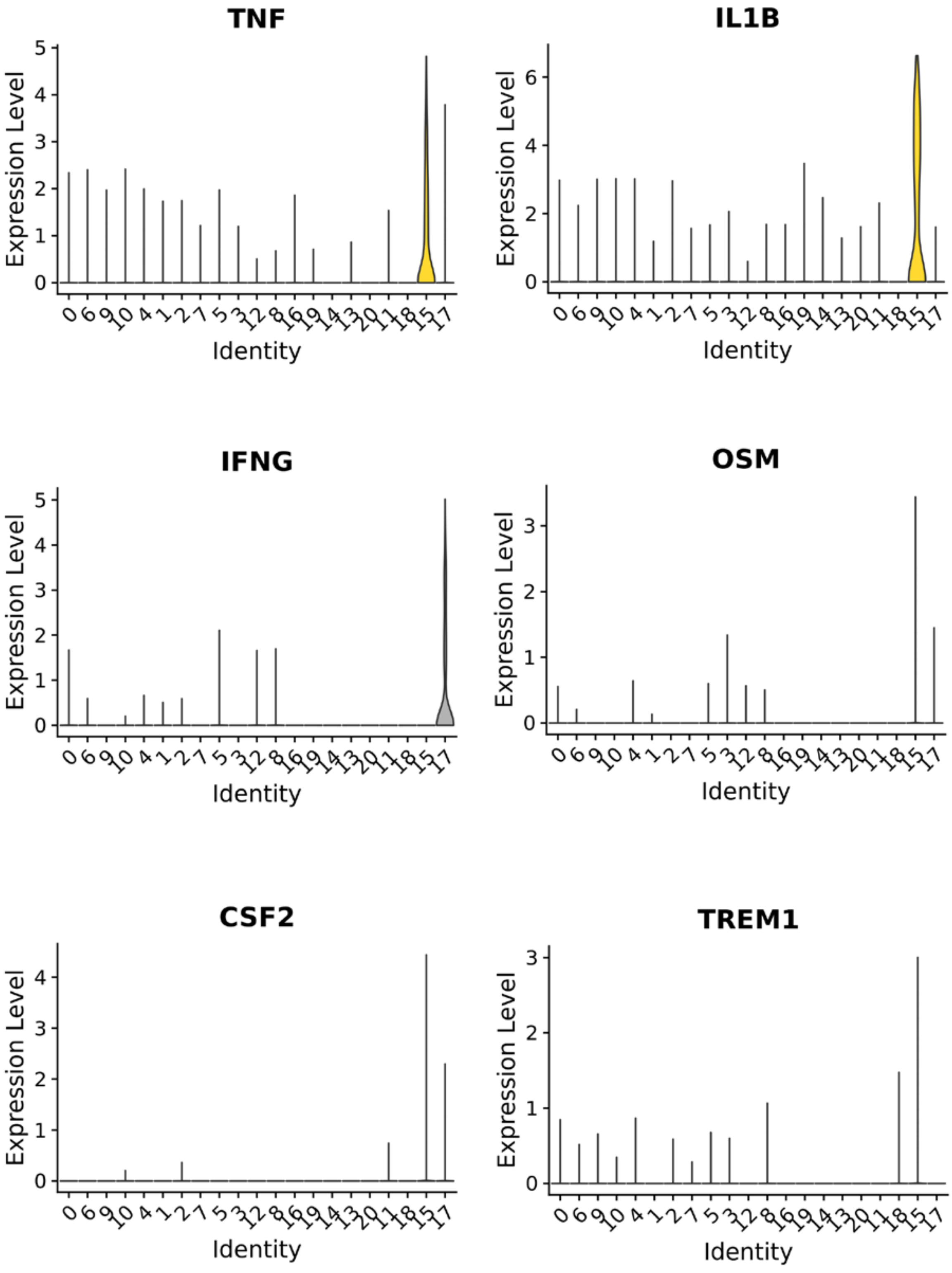

**Figure S16: Violin plots showing predominant expression of inflammatory regulators *TNF, IL1B, IFNγ, OSM, TREM1* and *CSF2* in clusters 15 and/or 17** (see also Figure 4, Table S5).

#### Figure S17

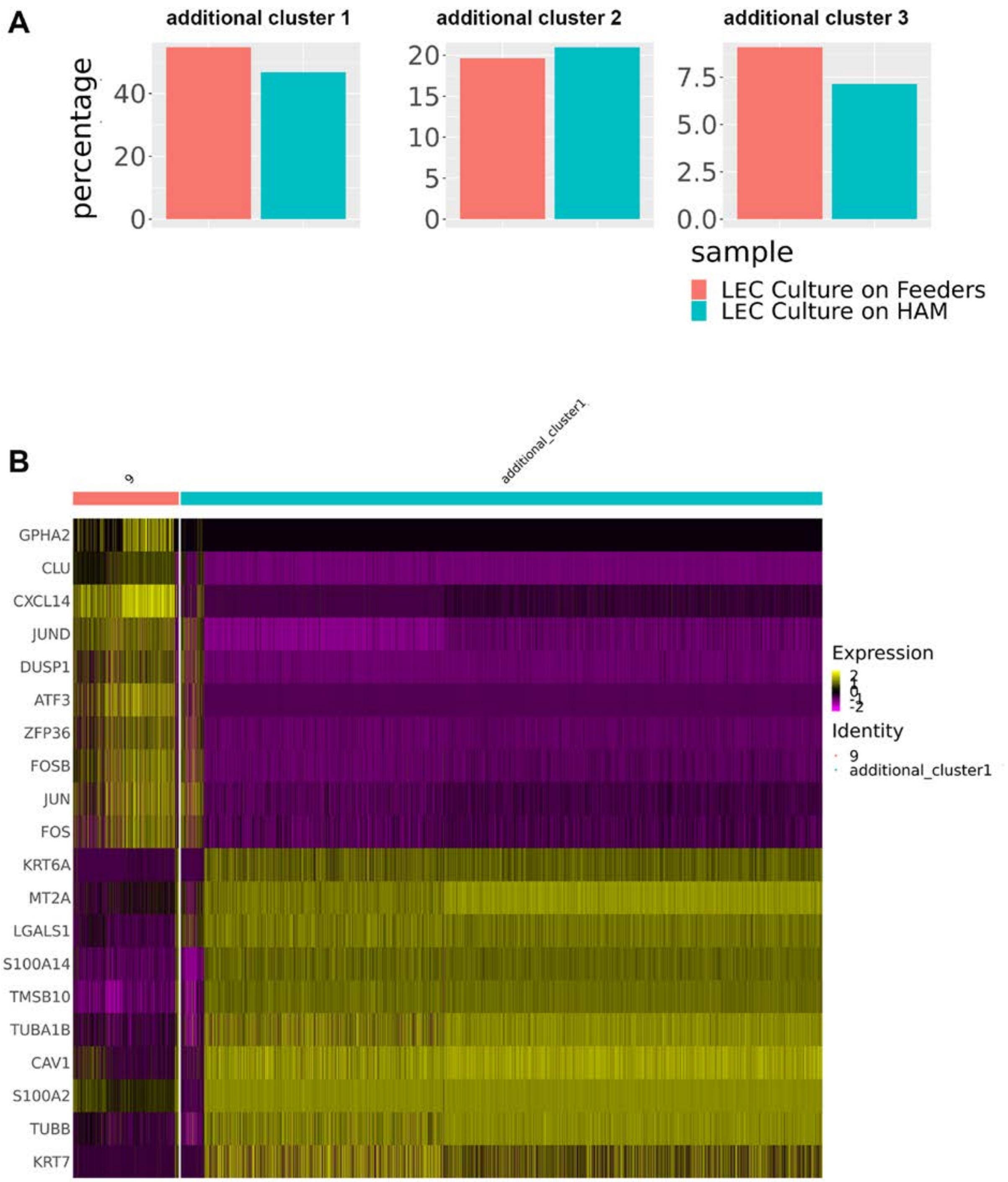

**Figure S17: scRNA-Seq of *ex vivo* expanded LECs and method comparison** (see also Figure 5 and Table S7). **A)** The expansion of limbal epithelial cells with 3T3 feeders or on human amniotic membrane (HAM) results in similar cultured cell populations. LECs – limbal epithelial cells; **B**) Comparative heatmap showing the downregulation of qLSCs markers and acquisition of proliferative limbal progenitor markers during the *ex vivo* expansion of LECs.

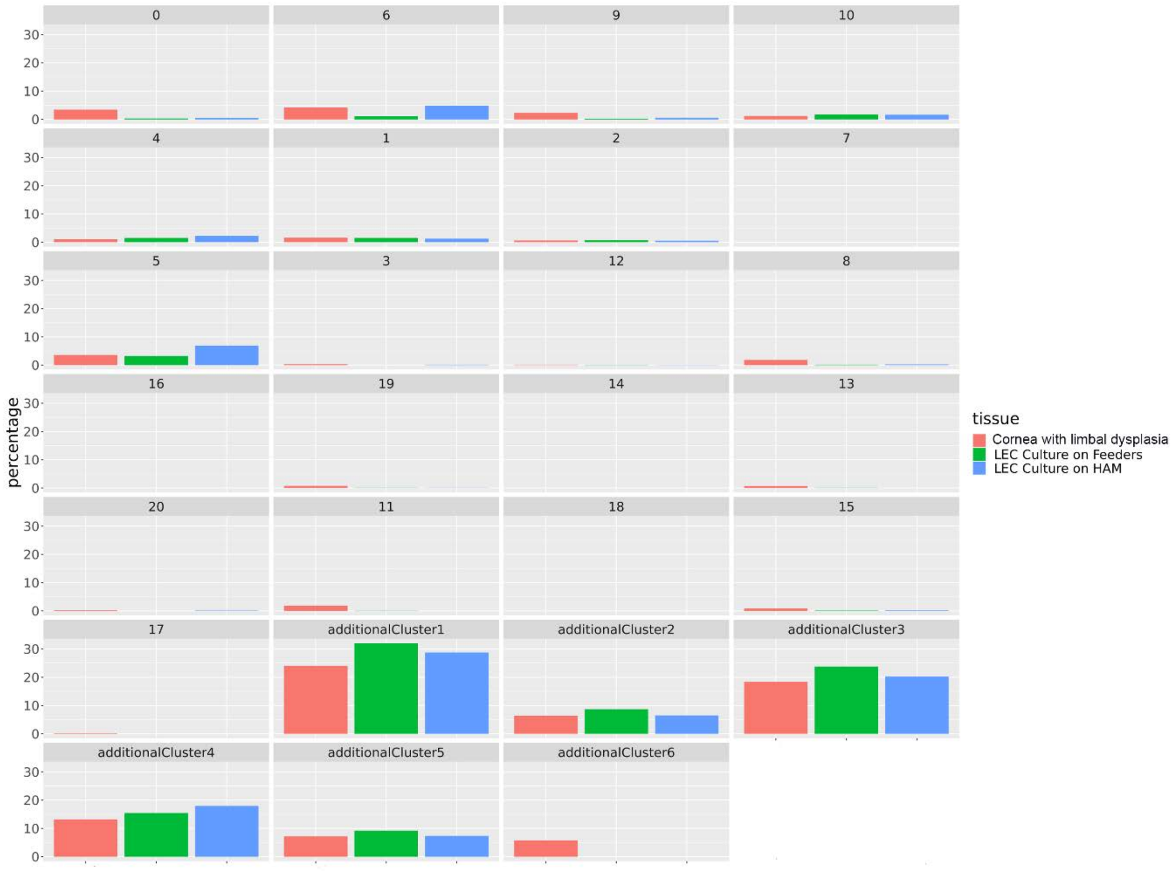

**Figure S18**

**Figure S18: scRNA-Seq analysis shows the presence of six additional clusters in the corneal sample with limbal dysplasia** (see also Figure 6, Figures S18, S19 and Table S9). The additional clusters 1-5 were also found in the *ex vivo* expanded LECs. LEC – limbal epithelial cell.

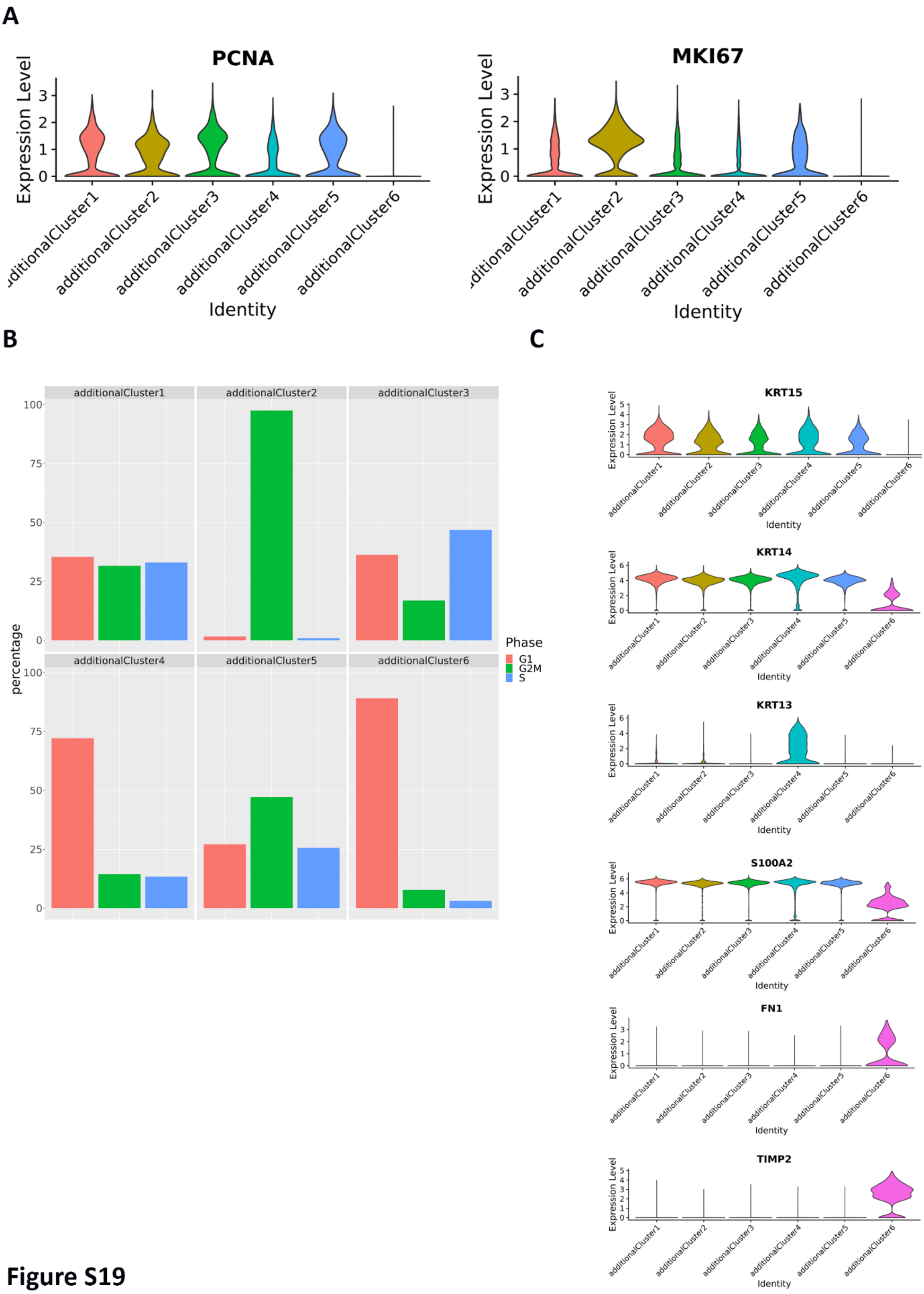
A
PCNA
MKI67
Expression Level
Identity
additionalCluster1
additionalCluster2
additionalCluster3
additionalCluster4
additionalCluster5
additionalCluster6
B
percentage
Phase
G1
G2M
S
C
KRT15
KRT14
KRT13
S100A2
FN1
TIMP2

**Figure S19**

**Figure S19: Detailed analysis of additional clusters 1-6 found in the cornea with limbal dysplasia** (see also Figure 6, Figures S18 and Table S9). **A**) High expression of proliferative markers *PCNA* and *MKi67* in the additional clusters 1-5; **B**) Cell cycle analysis of additional clusters 1-6; **C**) Violin plot showing expression of limbal basal epithelial (*KRT15, KRT14*), conjunctival (*KRT13*), proliferative limbal progenitor (*S100A2*) and fibroblasts markers (*FN1, TIMP2*).

**Figure S20**

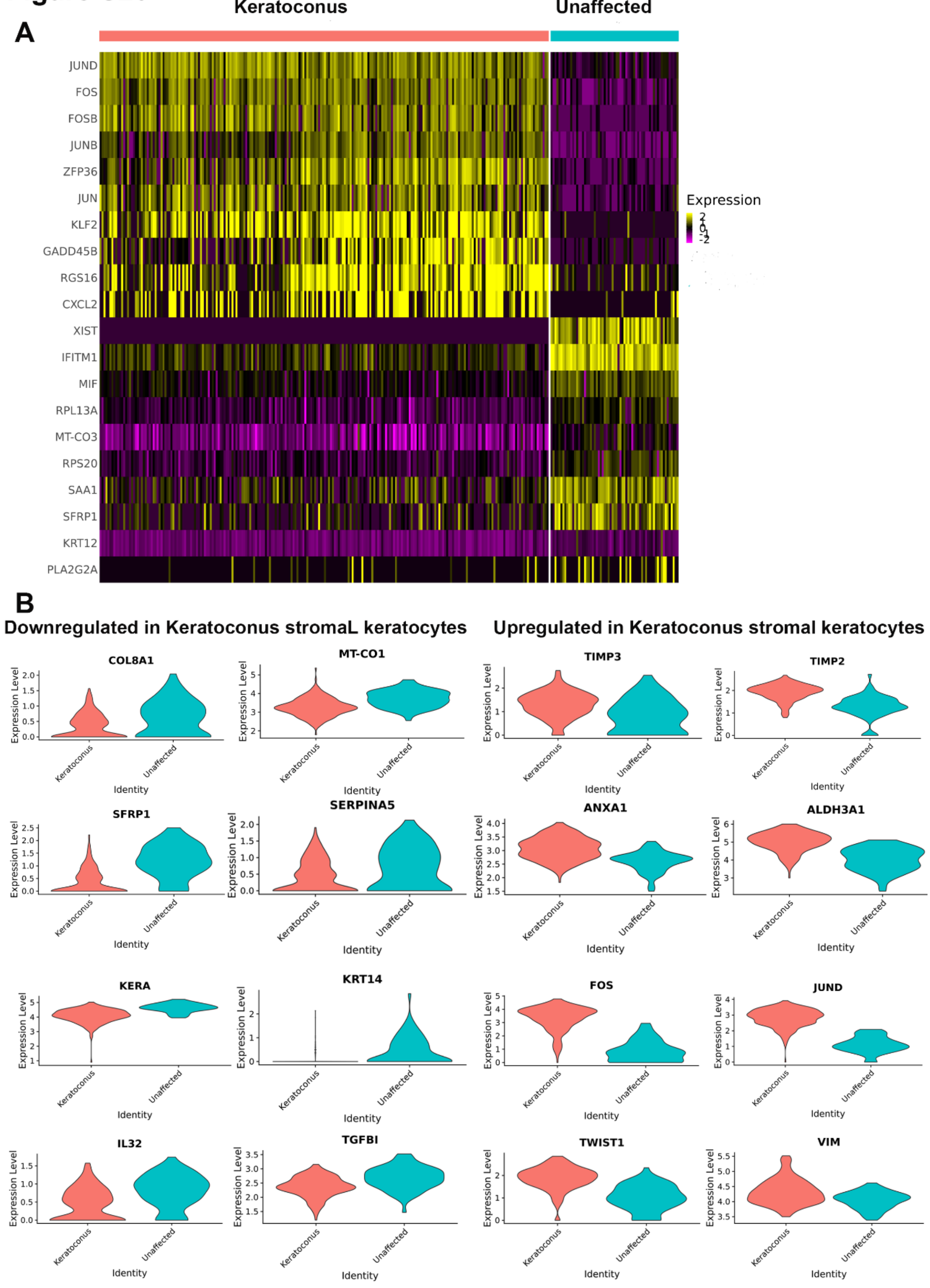

**Figure S20: Detailed characterisation of corneal stroma keratocytes in keratoconus cornea patients** (see also Figure 7, S21, Tables S10-13)**. A**) Comparative heatmap showing gene expression differences in corneal stroma keratocytes between the Keratoconus and unaffected subjects; **B**) Violin plots of key candidate genes that are either downregulated or upregulated in keratoconus stromal keratocytes.

#### Figure S21

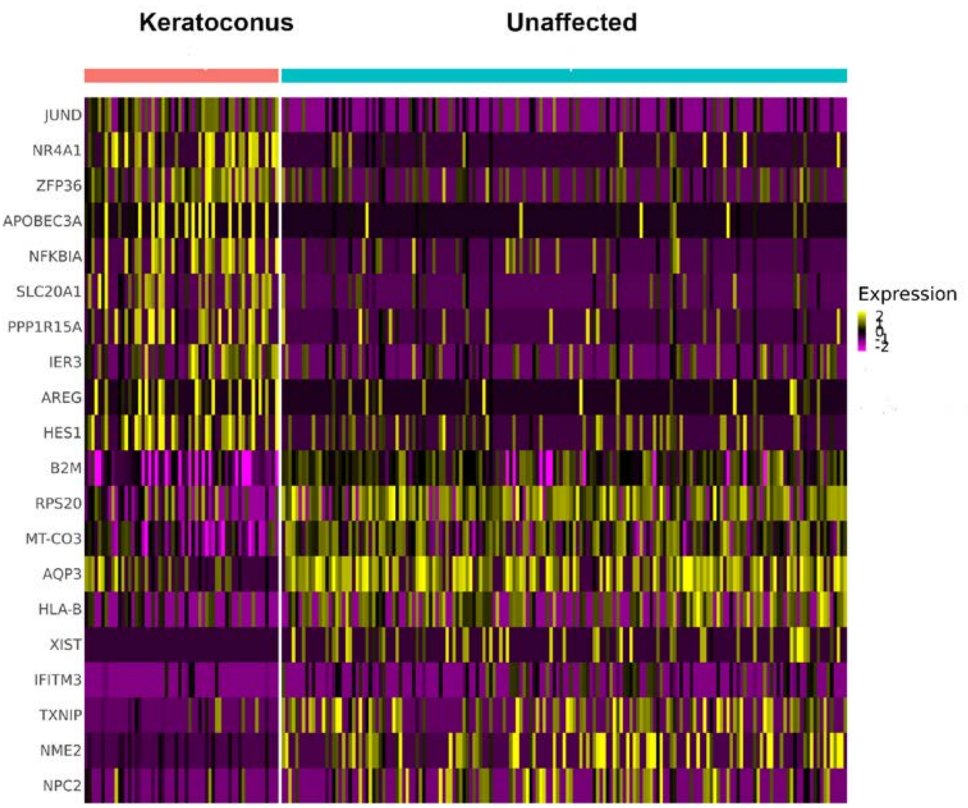

**Figure S21: Detailed characterisation of TAs in keratoconus patients** (see also Figure 7, S20, Tables S10-13)**. A**) Comparative heatmap showing gene expression differences in TAs (cluster 4) between the keratoconus and unaffected subjects.

#### Materials and Methods

##### Human Tissue Donation

Four adult human eyes (4: 51, 75, 83 and 86 years old), three central corneas and eight cornea-scleral rings were donated for research following informed consent. The adult eye tissue was provided by NHS Blood and Transplant Tissue and Eye Services following ethical approval (18/YH/04/20). The human fetal material was provided by the Joint MRC/Wellcome Trust (MR/R006237/1) Human Developmental Biology Resource (Gerrelli et al., 2015).

##### Single cell sequencing

Cornea-conjunctival tissue was excised from donated eyes and dissociated to single cells using a multi tissue dissociation kit (Miltenyi Biotec). Dissociation time varied from 15 to 45 minutes depending on the age of the tissue and a gentleMACS Dissociator (Miltenyi Biotec) was used to aid dissociation of the adult tissue. Cell count and viability were monitored using a Tali Image-Based Cytometer and Viability Kit (Thermo Fisher Scientific).

For scRNA-Seq cells were captured and libraries generated using the Chromium Single Cell 3′ Library & Gel Bead Kit, version 3 (10x Genomics). scRNA-Seq libraries were sequenced to 50,000 reads per cell on an Illumina NovaSeq 6000.

For scATAC-Seq a nuclei preparation from the dissociated cells was performed following recommendations from 10x Genomics. The subsequent nuclei were captured and sequencing libraries generated using the Chromium Single Cell ATAC Library & Gel Bead Kit, version 1 (10x Genomics). scATAC-Seq libraries were sequenced to 25,000 reads per nucleus on an Illumina NovaSeq 6000.

**Analysis of single cell sequencing**

The sequenced samples were de-multiplexed and aligned to human reference genome GRCh38 before being quantified using CellRanger version 3.01. Quality control filtering was applied to remove any cells where fewer than 1000 reads or 500 genes or greater than 15% mitochondrial reads. Diagnostic plots were used to determine the most appropriate thresholds for our data. DoubletFinder was used to predict doublets in the data which were then filtered.

The Seurat R package (version 3.1.3) was used to normalise individual experiments using the “LogNormalize” method. The Seurat standard integrated analysis approach was used to overcome batch effects and combine samples from the developmental and adult samples (Stuart et al., 2019). Firstly we selected a subset of 2000 genes which were highly variable genes, The “FindVariableFeatures”. We then chose the first 30 principle components were used for integration. A combined dataset was created by finding anchors between the individual datasets to create a batch corrected expression matrix. The corrected datasets were then clustered using a graph-based clustering. We used Clustree to assess the stability of clusters from a resolution of 0.2 - 1 and determined that a resolution 0.6 gave the highest number of stable clusters with cells from each donor represented in each cluster. The FindMarkers function identified markers for each cluster. Cell types were then assigned to these clusters and annotated using these genes lists. Cluster identity was validated using immunohistochemistry. The clustering results were visualised using uniform manifold approximation and projection (UMAP).

Integrated cluster analysis, described above, was performed on the following groups of developmental samples: 10PCW, 12PCW, 13-14PCW, 16PCW, 17-18PCW, 20-21 PCW. The adult dataset was used as a reference to predict cell identities in the developmental data using the Seurat “TransferData” function.

Cells within clusters 1, 2, 4, 5,7 and 9 from the adult datasets were selected for pseudotime analysis. The Harmony batch correction method, which uses soft clustering to overcome over-discretisation, was used to remove batch effects between donors. A trajectory and ordering of cells were inferred using Monocle 3.

The gene lists were analysed using the upstream regulator function from QIAGEN Ingenuity Pathway Analysis (QIAGEN IPA).  CellPhoneDB was used to predict interacting cells across clusters. The analysis was performed using the clusters identified by Seurat.

CellRanger ATAC (version 1.2) was used to identify accessible regions from scATAC-Seq sequencing files. The cellranger-atac agrr module was used to create a set of shared peaks present across all four samples. The samples were then analysed with Signac version 0.2.4 along with Seurat. The percentage of reads in peaks, number of reads in percentage of reads in peaks per cell, percentage mapping to ENCODE blacklist, nucleosome signal and TSS enrichment was calculated. The data was filtered using thresholds to remove cells where the number of fragments within peaks was less than 1000 or greater than 50000, or with a blacklist ration of greater than 5% or a nucleosome signal of greater than 10 and a TTS enrichment of greater than 2. A gene activity matrix was calculated in the scATAC-Seq dataset, using the closest genes to the peaks. Seurat was then used to perform clustering analysis on each sample and transfer the cell type labels from the scRNA-Seq to the scATAC-Seq data. Seurat integration methods were applied to the data peak accessibility to overcome donor effects and combine the four ATAC-Seq datasets. Signac was used to create “pseudo-bulk” accessibility tracks by grouping cells by cell type within promoter/enhancer regions of key genes.

**Adult and developmental human cornea and conjunctiva immunohistochemistry**

For immunohistochemistry (IHC) adult and developmental eyes were fixed in 4% PFA overnight followed by three washes in phosphate-buffered saline (PBS), incubated overnight in 30% sucrose/PBS, embedded in optimum cutting temperature (OCT) embedding matrix

(Cellpath) and frozen at -20°C. 10 µm cryostat sections were collected using a Leica Cm1860 cryostat (Leica). Cryosections were air-dried, washed several times in PBS and incubated in blocking solution (10 % normal goat serum, 0.3 % Triton-X-100 in PBS) for one hour at room temperature. Slides were incubated with the appropriate primary antibody overnight 4°C (**Table S15**). After rinsing with PBS, sections were incubated with the secondary antibody for 2 hours at RT. Alexa Fluor 488 and 546 secondary antibodies (Invitrogen-Molecular Probes) were used at a 1:1000 dilution. Negative controls were carried out by omitting the primary antibody. Afterwards, sections were washed three times in PBS and mounted with Vectashield (Vector Laboratories) containing 10 µg/ml Hoechst 33342 (Life Technologies) for counterstaining nuclei. All details of antibody purchase and concentrations are provided in **Table S15**.

**Human Limbal Explant Culture**

Adult human limbal tissue was obtained from the surplus cornea-scleral rings remaining from the cornea-scleral buttons provided by NHSBT for full thickness corneal grafts (penetrating keratoplasty) procedures and an exenterated globe provided by NuTH Hospitals NHS Foundation Trust. The human amniotic membrane (HAM) was obtained from placentas donated during elective caesarean section deliveries and supplied by NHSBT. HAM is processed, mounted on nitrocellulose paper and frozen at -80$^{o}$C in 50% glycerol/Hanks solution and supplied as individual 3cm$^{2}$ units. In accordance with the Declaration of Helsinki, informed consent for the use of human tissue for experimental research was obtained.

LEC culture was performed by preparing the epithelial medium, HAM construct and limbal biopsy preparation and placement onto the HAM. Epithelial medium was prepared to the following composition: 66.75% Low Glucose Dulbecco's Modified Eagle Medium (DMEM), 22.25% Ham's F12 medium (Gibco), Human Serum AB 10%, hydrocortisone 0.4µg/ml, insulin 5µg/ml, triiodothyronine 1.4ng/nl, adenine 24µg/ml, cholera toxin 8.4ng/ml and EGF 10ng/ml (Sigma Aldrich), then sterile filtered via 0.2µ filter (ThermoFisher Scientific). Three separate aliquots of the resultant media were further supplemented with 1% penicillin/streptomycin (Gibco). The HAM was defrosted at room temperature and in a laminar air hood washed twice by submersing the tissue in Dulbecco's Phosphate Buffered Saline (DPBS), (Gibco), supplemented with 1% penicillin/streptomycin. Subsequently, the tissue was washed a third time in 1% penicillin/streptomycin supplemented epithelial medium. Two 24mm$^{2}$ glass coverslips were prepared as per the HAM washing procedure, in separate wells to the HAM. One glass coverslip was carefully placed onto the lid of a sterile 6 well plate using sterile forceps and a drop of 1% penicillin/streptomycin supplemented epithelial medium added.

The HAM was carefully separated from the nitrocellulose paper with sterile forceps and stretched over the coverslip, avoiding air bubbles and ensuring the epithelial side of the HAM was facing up, verified by brief application of a cellulose eye spear to the HAM. The overhanging edges of the HAM were trimmed using a sterile scalpel leaving approximately 1mm overlap on each side. The second glass coverslip was placed on the edge of the 6 well plate lid and a drop of media added. The HAM/coverslip was carefully lifted using the edge of a sterile scalpel blade and sterile forceps then the edges were tucked under the coverslip. The HAM was secured by placement on top of the second coverslip, avoiding air bubbles between the coverslips and ensuring that the HAM remains flat and *in situ* throughout the culture process. The HAM construct was placed into a well of a fresh sterile 6 well plate with a drop of media underneath the construct, preventing air bubbles. Finally, the HAM was covered in the epithelial culture medium supplemented with 1% penicillin/streptomycin.

The limbal biopsies were prepared from the human cornea-scleral rings, dissected to $2mm^2$ consisting of approximately $1mm^2$ peripheral cornea and $1mm^2$ adjacent cornea, ensuring all the cornea-scleral limbus was included. Due to the manual dissection, small rims of surrounding conjunctiva were also present in the dissected limbal rings. The stromal side remained in place. Each biopsy was placed at the centre of the prepared HAM, the spent epithelial medium removed prior to biopsy attachment. After an interlude of 2 minutes, 1% penicillin/streptomycin supplemented epithelial medium with was added slowly to each culture to ensure the explant was covered in medium without causing it to detach from the HAM. The cultures were fed with antibiotic-free epithelial medium on the third day and then every other day thereafter. Cultures were terminated when outgrowth reached approximately 90% confluence over the HAM. All cultures were performed under identical conditions and incubated in a tissue culture incubator at $37^{O}C$ humidified with 5% $CO_2$

**Human limbal epithelial cell culture on 3T3-J2 feeder layers**

Cadaveric adult human limbal tissue was obtained from the cornea-scleral rings remaining after removal of the central cornea for transplantation supplied by the NHS Blood and Transplant (NHSBT) Tissue and Eye Services. Human tissue was handled according to the tenets of the Declaration of Helsinki and informed consent was obtained for research use of all human tissue from the next of kin of all deceased donors. The study was approved by the NRES Committee North East - Newcastle & North Tyneside 1 (REC number: 18/YH/04/20).

Twenty-four hours before LEC isolation from cornea-scleral tissue, mitotically inactivated J2–3T3 mouse fibroblasts were suspended in high-glucose DMEM supplemented with bovine calf serum (10%) (Hyclone, USA) and penicillin/streptomycin (1%) (Thermo Fisher Scientific, USA) and plated in a 9.6 $cm^2$ tissue culture well at the final density of $2.4 \times 10^4$ cells per $cm^2$ as previously described (Yu et al., 2016). The use of bovine calf serum instead of fetal calf serum was recommended by the manufacturer of the 3T3-J2 cell line (Karafast, USA). The 3T3 cell suspension was placed in a tissue culture incubator at 37°C overnight to allow the establishment of a 3T3 feeder layer. On the following day, LECs were harvested from cadaveric cornea-scleral rims as previously described (Ahmad et al., 2007). The deeper layers of the cornea-scleral rings were dissected away together with excess sclera leaving a ring containing approximately 2 mm of peripheral cornea and 2 mm of adjacent conjunctiva. The remaining tissue containing limbal epithelium was then cut into smaller 1 $mm^2$ pieces. The LECs were isolated from these pieces using serial trypsinization with 0.05% trypsin-EDTA solution (Thermo Fisher Scientific, USA). After 20 minutes incubation in a tissue culture incubator, the resulting cell suspension was removed from the limbal pieces and epithelial medium was added to this suspension. After the cell suspension was centrifuged for 3 minutes at 1000 rpm in Heraeus Megafuge 16R Centrifuge (Thermo Fisher Scientific, USA), the supernatant was removed and the remaining cell pellet was re-suspended in epithelial medium containing 3:1 mixture of low-glucose DMEM:F12 supplemented with fetal calf serum 10%, penicillin/streptomycin 1% (all Thermo Fisher Scientific, USA), hydrocortisone 0.4 µg/ml, insulin 5µg/ml, triiodothyronine 1.4 ng/ml, adenine 24 µg/ml, cholera toxin 8.4 ng/ml and EGF 10 ng/ml (all Sigma-Aldrich, UK). The trypsinization and centrifugation process was repeated a further three times using the same limbal tissue and the same centrifuge settings. The resulting cell suspensions were pooled together. Cells were counted and assessed for viability using trypan blue exclusion and a haemocytometer. 30,000 viable LECs in epithelial medium were added to one 9.6 $cm^2$ tissue culture well containing the growth arrested 3T3 fibroblast and placed in a tissue culture incubator at 37°C with a humidified atmosphere containing 5% $CO_2$. The medium was exchanged on the third culture day and every other day thereafter. Several days after, cell colonies with typical morphology started to appear and were cultured until they became sub-confluent. After 3T3 feeder cells were detached and removed using 0.02% EDTA (Lonza, Switzerland), sub-confluent primary cultures were dissociated with 0.5% trypsin-EDTA (Santa Cruz, USA) to single cell suspension and passaged at a density of $6 \times 10^3$ cells/$cm^2$. For serial propagation, cells were passaged and cultured as above, always at the stage of sub-confluence, until they reached passage 3.

**Human LEC siRNA Transfection**

Passage one human LECs from 3 different donors were grown on 3T3 feeder layer in complete epithelial medium supplemented with EGF, adenine, cholera toxin, hydrocortisone, insulin and triiodothyronine. A day before transfection, LECs ($150x10^3$) were re-seeded in 12-well plate without feeders in order to increase transfection efficiency. The day after re-seeding cells were transfected with either *GPHA2 (*HSS153169), or *TFPI2* (HSS111884) Human Stealth siRNAs and Stealth RNAi siRNA Negative Control Lo GC using Lipofectamine™ RNAiMAX Transfection Reagent (ThermoFisher Scientific) according to the manufacturer's protocol. After 48 hours incubation with siRNA, cells we re-seeded into 6 well plates for colony forming efficiency and clonal assays and cultured on 3T3 feeders for 14 days. The rest of the cells was used for qRT-PCR.

**Colony forming efficiency (CFE) assay**

Mitotically inactivated 3T3-J2 mouse embryonic fibroblasts (Kerafast, USA) were suspended in complete medium containing: high-glucose DMEM (89%), FBS (10%) and penicillin/streptomycin (1%) and plated in a 9.6 $cm^2$ tissue culture well at a final density of $2.4×10^4$ cells per $cm^2$ and placed in a tissue culture incubator overnight to allow the establishment of a 3T3 feeder layer. The following day, 500 viable LECs were plated onto the prepared 3T3 feeder cells together with 2 ml of epithelial medium. The CFE culture was then placed in the tissue culture incubator and the epithelial medium was changed on the third day and then every second day thereafter with regular microscopic examination (Eclipse TS100, Nikon, Japan) for the presence of colonies. The CFE was measured on the 12th day of the culture. This was performed by removal of the epithelial medium followed by two brief washes with PBS. The culture was then fixed with 3.7% formaldehyde (VWR International, UK) in PBS for 10 minutes at room temperature. Next, the formaldehyde solution was removed, and the culture was irrigated with PBS. The colonies were then stained by incubation with 1% Rhodamine B (Sigma-Aldrich) in methanol for 10 minutes at room temperature. Following staining, the colonies were counted under dissecting microscope (SMZ645, Nikon, Japan). The CFE was calculated using the formula: number of colonies formed/number of cells plated ×100.

**Clonal assays**

The clonal type was determined by (1) the morphology of colonies and (2) the percentage of aborted colonies as follows: when <5% of the total colonies were terminally differentiated, the clone was scored as a holoclone; when more than 95% of colonies were terminally differentiated, the clone was scored as a paraclone and finally, when >5% but <95% of colonies were terminally differentiated, the clone was classified as a meroclone (Barrandon 1989 and Pellegrini 1999).

**Immunocytochemistry of LECs**

Cultured LECs were fixed for 15 minutes in 4% PFA. A blocking step was performed by incubation in antibody diluent containing 1% bovine serum albumin (Sigma-Aldrich) with 5% normal goat serum (Thermo Fisher Scientific) for 30 minutes prior to staining. Permeabilization with 0.2% Triton X-100 in PBS was performed prior to staining with antibodies for internal cell markers. Cells were incubated with the appropriate primary antibodies (**Table S15**) at 4°C overnight and further incubated with secondary antibodies for 1 hour at room temperature. Following this, cells were washed and then mounted in Vectashield anti-fading media containing Hoechst (Vector Laboratories, UK). All details of antibody providers and concentrations are provided in **Table S15**.

**Image acquisition and processing**

Adult and developmental eye sections and cultured LECs were viewed on a Zeiss Axio ImagerZ2 equipped with Apotome 2 and Zen 2012 blue software (Zeiss, Germany). Objectives lens used were EC Plan Neofluar 20x/0.5 Ph2 and EC Plan Apochromat 63x/1.4 Ph3. Series of XZ optical sections (<1 µm thick) were taken at 1.0 µm steps throughout the depth of the section. Final images are presented as a maximum projection and adjusted for brightness and contrast in Adobe Photoshop CS6 (Adobe Systems).

**Quantitative Reverse Transcriptase Polymerase Chain Reaction (qRT- PCR)**

Following siRNA treatment RNA was extracted from cultured LECs using the ReliaPrep RNA Cell Miniprep System (Promega). cDNA was then synthesised using the GoScript Reverse Transcription System as per the manufacturer's protocol. qPCR was then performed using Go-Taq qPCR Master Mix (Promega) and was composed of 5 µl GoTaq, 0.5 µl forward primer, 0.5 µl reverse primer, 0.5 µl template cDNA, 3.4 µl RNAse-free water and 0.1 µl CXR. All reactions were analysed on a QuantStudio™ 7 Flex Real Time PCR System (ThermoFisher Scientific) according to the manufacturer's instructions. A standard, 40-cycle qPCR was performed for each sample.  The primer sequences used for qRT-PCR are listed in **Table S16**. The data was analysed using the $2^{-\Delta\Delta Ct}$ method.

**Data availability**

All single cell data are deposited in the Gene Expression Omnibus.
